# Comparative analysis of CPI-motif regulation of biochemical functions of actin capping protein

**DOI:** 10.1101/2020.02.06.936211

**Authors:** Patrick McConnell, Marlene Mekel, Alexander G. Kozlov, Olivia L. Mooren, Timothy M. Lohman, John A. Cooper

## Abstract

The heterodimeric actin capping protein (CP) is regulated by a set of proteins that contain CP-interacting (CPI) motifs. Outside of the CPI motif, the sequences of these proteins are unrelated and distinct. The CPI motif and surrounding sequences are conserved within a given protein family, when compared to those of other CPI-motif protein families. Using biochemical assays with purified proteins, we compared the ability of CPI-motif-containing peptides from different protein families to a) bind to CP, b) allosterically inhibit barbed-end capping by CP, and c) allosterically inhibit interaction of CP with V-1, another regulator of CP. We found large differences in potency among the different CPI-motif-containing peptides, and the different functional assays showed different orders of potency. These biochemical differences among the CPI-motif peptides presumably reflect interactions between CP and CPI-motif peptides involving amino-acid residues that are conserved but are not part of the strictly defined consensus, as it was originally identified in comparisons of sequences of CPI motifs(*1, 2*) across all protein families (*1, 2*). These biochemical differences may be important for conserved distinct functions of CPI-motif protein families in cells with respect to the regulation of CP activity and actin assembly near membranes.

## INTRODUCTION

The assembly of actin filaments is a key element of how cells control their shape, move about, and exert force. Actin filaments grow and shrink by the addition and loss of subunits from their ends, both barbed and pointed (*3–6*). The dynamics of subunit addition and loss at barbed ends of actin filaments is controlled by a number of proteins, including the heterodimeric alpha/beta actin capping protein (CP), which is nearly ubiquitous among eukaryotic cells and tissues.

In cells, CP is negatively modulated by direct interactions with the protein V-1 / myotrophin and by proteins with CP-interacting (CPI) motifs (*1*). Inhibition by V-1 occurs by a steric blocking mechanism in which V-1 directly competes with F-actin barbed ends for binding to CP (*7*). Inhibition by CPI-motif proteins occurs via an allosteric mechanism, and CPI-motif binding decreases the affinity of CP for both F-actin barbed ends and V-1 (*8*). Cell cytoplasm contains relatively high concentrations of V-1 and CP, and both proteins appear to diffuse freely about the cell (*9, 10*). In contrast, CPI-motif proteins are present in far lower amounts, and they are generally targeted to membranes (*11–13*). Proteins with CPI motifs are otherwise unrelated to each other, and cells may express one or more CPI-motif proteins (reviewed in (*11, 14*)). The different proteins are generally targeted to membranes by other domains of the proteins, where they are proposed to promote actin polymerization and actin-based motility(*11, 14*).

These observations led to a model, proposed by Fujiwara and colleagues (*8*), in which V-1 inhibits CP globally, and CPI-motif proteins activate CP locally by inducing dissociation of V-1. This model(*8*) is supported by several studies in cells. First, V-1 has been shown to inhibit CP in cells (*9*). Second, a number of CPI-motif proteins have been found to require interaction with CP for their cellular activity (reviewed in (*11, 14*)). Third, mutations of CP that inhibit the binding of CPI-motif proteins lead to an apparent loss of function of CP in cells (*15*). While this model and its supporting evidence are compelling, in our view, the potential validity of the model does not exclude other potential models of functions for CPI-motif proteins in cells.

To be specific, the functions of CPI-motif proteins in cells have been proposed to include the following set of activities, which we emphasize are not exclusive of one another. These proposals have been made based on biochemical and cellular studies, notably the allosteric nature of the effect of CPI-motif proteins on the ability of CP to bind to barbed ends of actin filaments and to bind V-1. The proposed functions include the following: 1) targeting of CP to a membrane location where CP binds and caps actin filament barbed ends, 2) inducing the dissociation of CP (uncapping) from a capped barbed end, thereby allowing that barbed end to shrink or grow, 3) inducing the dissociation of V-1 from CP, converting CP from a capping-inactive to a capping-active form. We consider the possibility that a given CPI-motif protein may exhibit one or more of these functions, with greater or lesser potency, relative to other CPI-motif proteins.

To investigate the biochemical basis for how the different types of CPI-motif proteins may function in cells, we examined the interactions among CP, actin barbed ends, V-1 and CPI-motifs, using purified components in vitro. Many of these interactions have been studied in previous work, in studies of one or more of these components and one or more of the various CPI-motif proteins (*1, 2, 7, 8, 11, 14, 16–19*). Our goal here was to provide a comparative analysis of the different CPI-motif proteins, one with another, and examine how their effects on actin capping and V-1 binding correlate with each other and with their affinity for binding CP.

## MATERIALS AND METHODS

### Proteins and Peptides

His-tagged mouse non-muscle F-actin-capping protein alpha-1 subunit (CPα1, Q5RKN9, Mus musculus) and beta-2 subunit (CPβ2, Q923G3, Mus musculus) were co-expressed in bacteria and purified as described (*20*). Myotrophin (V-1, P58546, Homo sapiens) was expressed and purified as described (*20*). N-carboxytetramethylrhodamine (TAMRA)-V-1 (C45S, C83S) was prepared as described (*21*). Rabbit skeletal muscle alpha-actin (actin, P68135, Oryctolagus cuniculus) was prepared as described (*22*). The concentrations of purified proteins were determined by UV absorbance with extinction coefficients as follows: CPα1β2 ε_280nm_ = 78310 M^-1^ cm^-1^, V-1 ε_280nm_ = 9970 M^-1^ cm^-1^, actin ε_290nm_ = 26460 M^-1^ cm^-1^.

CPI-motif peptides used in this study were designed based on crystal structures of CP in complex with CPI-motif regions of CARMIL1 and CD2AP (*1*). The peptide boundaries were chosen so that a) the peptides were the same length, b) the peptides included and were centered around the CPI motif, and c) the CARMIL-derived peptides did not include the CSI motif. The peptides were derived as follows, with peptide boundaries and UniProtKB designations for the proteins: Capping protein Arp2/3 Myosin-I Linker (CARMIL1 G969-A1005, Q5VZK9, Homo sapiens), Capping protein, Arp2/3 and Myosin-I linker protein 3 (CARMIL3 E959-M994, Q8ND23, Homo sapiens), WASH complex subunit 2C (WASHCAP A990-R1026, Q9Y4E1, Homo sapiens), CapZ-interacting protein (CapZIP V140-R176, Q6JBY9, Homo sapiens), CD2-associated protein (CD2AP D473-H509, Q9Y5K6, Homo sapiens), SH3 domain-containing kinase-binding protein 1 (CIN85 L463-S499, Q96B97, Homo sapiens), Casein kinase 2-interacting protein-1 (Pleckstrin homology domain-containing family O member 1, CKIP-1/PLEKHO R137-M173, Q53GL0, Homo sapiens), and Twinfilin-1 (Twf-1 H317-D350, Q91YR1, Mus musculus). The Twf-1 peptide was identical to one previously described (*21*). The CPI-motif peptide from human CARMIL2 was tested, but it had poor solution properties and was not included in this study. As a negative control peptide, we used a mutated form of the CARMIL1 peptide with two amino acid substitutions (G969-A1005 / K987A R989A). This mutation was designed from structural and functional studies (*1*) and was shown to have very little CP-binding activity in biochemical assays (*17*).

Unlabeled (N-acetyl, C-amide) and labeled [(N-TAMRA), C-amide] CPI-motif peptides were purchased from WatsonBio Sciences (Houston, TX). Peptide concentrations were determined by absorbance using an Agilent Cary 100 UV-Vis spectrophotometer. TAMRA-labeled CPI-motif peptide concentrations were determined by absorbance at 554 nm (TAMRA ε_554nm_ = 80000 M^-1^ cm^-1^). Unlabeled CPI-motif peptides were dissolved in water and protein concentrations were determined by absorbance at 205 nm in PBS pH 7.4 with 0.005 % TWEEN 20 with extinction coefficients as follows: CARMIL3 ε_205nm_ = 123160 M^-1^ cm^-1^, WASHCAP ε_205nm_ = 126360 M^-1^ cm^-1^, CARMIL1 ε_205nm_ = 127470 M^-1^ cm^-1^, CapZIP ε_205nm_ = 140850 M^-1^ cm^-1^, CD2AP ε_205nm_ = 136510 M^-1^ cm^-1^, CIN85 ε_205nm_ = 118330 M^-1^ cm^-1^, CKIP-1 ε_205nm_ = 132090 M^-1^ cm^-1^, Twf-1 ε_205nm_ =116540 M^-1^ cm^-1^ (*23*). Peptide concentrations were confirmed by SYPRO Orange staining of SDS-polyacrylamide gels.

### Multiple Sequence Alignments and Phylogenetic Tree

Multiple sequence alignment of CPI-motif peptides was performed using the ClustalW algorithm within the DNASTAR Lasergene Suite/MegAlign Pro application (MegAlign Pro 15.0, DNASTAR, Madison, WI). An unrooted phylogenetic tree was generated from the aligned CPI-motif peptides using Interactive Tree of Life (iTOL V4.4.2, https://itol.embl.de/, (*24*)).

### Fluorescence Titration Binding Experiments

Fluorescence intensity and anisotropy titration experiments were performed on a one-channel L-format QM-2000 spectrofluorometer with FelixGX software (Photon Technology International/HORIBA Scientific, Piscataway, NJ) with excitation at 552 nm and emission detected at 582 nm. For intensity experiments, CP was titrated into 2 mL of 10 nM TAMRA-V-1 with and without 50 *µ*M CPI-motif peptide, with a 2-minute incubation after each addition. Based on binding affinities of the CPI-motif peptides for CP, a single-site binding model indicates that at concentrations of CP in the assay, the CPI-motif peptides are saturated at 20 *µ*M. For anisotropy experiments, CP was titrated into 2 mL of 20 nM TAMRA-CPI-motif peptide with and without 50 *µ*M V-1, with a 2-minute incubation after each addition. (20 *µ*M V-1 was used in one of the two CARMIL1 G969-A1005 titrations.) Based on the binding affinity of V-1 for CP, a single-site binding model indicates that at concentrations of CP in the assay, V-1 is saturated at 20 *µ*M. For competition experiments, unlabeled CPI-motif peptide was titrated into 2 mL of 10 nM or 20 nM TAMRA-labeled CPI-motif peptide in a preformed complex with CP at a concentration producing ∼80% saturation.

Experiments were performed at 25°C in 20 mM 3-(N-morpholino) propanesulfonic acid (MOPS), 100 mM KCl, 1 mM TCEP, 1 mM NaN_3_, 0.005 % TWEEN 20, pH 7.2. When salt and pH were varied, the conditions were 20 mM buffer, 100 to 300 mM KCl, 1 mM TCEP, 1 mM NaN3, 0.005 % TWEEN 20, and the pH buffers were as follows: MES pH 6.0, PIPES pH 6.5, MOPS pH 7.2, HEPES pH 7.5, EPPS pH 8.0. For each experimental value, one measurement was recorded every five seconds and averaged over 120 seconds. Equilibrium dissociation constants (K_d_) were determined from experiments in duplicate fit to a single-site binding model using MicroMath Scientist software (St. Louis, MO).

### Isothermal Calorimetry (ITC)

ITC experiments were performed on a MicroCal microcalorimeter (Malvern Panalytical, Malvern, PA). 2 *µ*M CP was titrated with 22 *µ*M of a given CPI-motif peptide at 25°C in 20 mM MOPS, 100 mM KCl, 1 mM TCEP, 1 mM NaN_3_, 0.005 % TWEEN 20, pH 7.2. For Twf-1, 10 *µ*M CP was titrated with 110 *µ*M of the CPI-motif peptide. For V-1, 4 *µ*M CP was titrated with 44 *µ*M V-1. Thermodynamic parameters were fit to a single-site binding model using MicroCal ITC Origin analysis software. Standard Gibbs free energy change was calculated using ΔG^0^ = - RTlnKa.

### V-1 Dissociation Rates Measured by Stopped-flow Fluorescence

V-1 dissociation experiments were performed on a SX.18MV stopped-flow instrument with Pro-Data SX software (Applied Photophysics Ltd., Leatherhead, UK). The excitation wavelength was 505 nm, and changes in emission intensity were detected using a 570-nm band-pass filter. 50 nM TAMRA-V-1 was pre-incubated with 250 nM CP. At time zero, the CP:TAMRA-V-1 complex was mixed via stopped flow with an equal volume of a solution containing 400 nM unlabeled V-1 plus a range of concentrations of CPI-motif peptide from 0-15 *µ*M. Experiments were performed at 25°C in 20 mM MOPS, 1 mM TCEP, 100 mM KCl, 1 mM NaN3, 0.005% Tween 20, pH 7.2. For every concentration of CPI-motif peptide, the mixing was repeated in replicates of 5-10, and traces were averaged. Apparent dissociation rates were determined by fitting the averaged data to a double-exponential decay model using Micromath Scientist software. The slower second step was much smaller in amplitude, did not correspond to photobleaching, and did not depend on the concentration of CPI-motif peptide in any consistent manner. Thus, every curve was fit to a double-exponential decay model, and only the result for the faster first step was used for further analysis. To determine the V-1 dissociation rate constant, the apparent dissociation rates as a function of CPI-motif peptide concentration were fit to a kinetic model, presented below, using Micromath Scientist software. The spontaneous dissociation rate constant for V-1 was calculated as the average of apparent dissociation rates in the absence of CPI-motif peptide from nine experiments.

#### Analysis of the kinetics of V-1 dissociation from capping protein V-1 complex (CP:V-1) upon addition of CPI-motif peptides

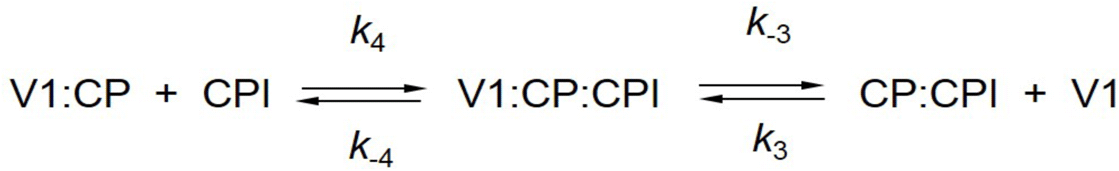

We used this kinetic scheme, derived from Steps 3 and 4 of the schemes shown in Figure 11. The dissociation of fluorescently labeled V-1 from CP, upon addition of an excess of CPI-motif

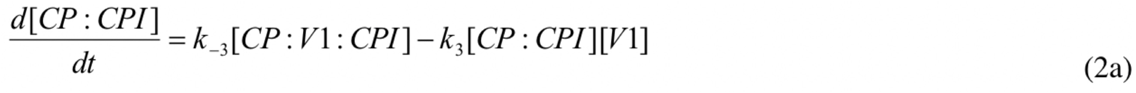

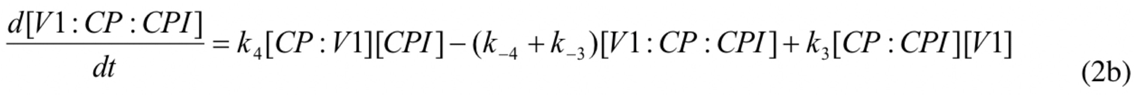

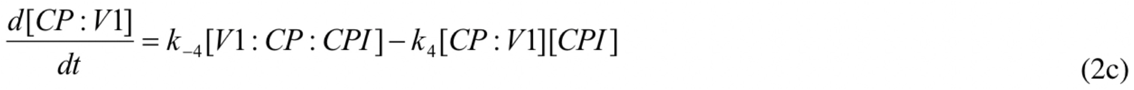

peptide, occurs via formation of a ternary complex intermediate V-1:CP:CPI. The differential equations describing the time dependences of the concentrations of capping protein complexes in the scheme are given in Equations 2a - 2c.

Under pseudo first order conditions ([*V-1*],[CPI]>[CP]) the concentrations of *V-1* and *CPI* can be treated as constant and the above set of differential equations (Eq 2) reduces to the first-order linear homogeneous system of differential equations written below in matrix form as 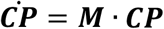.

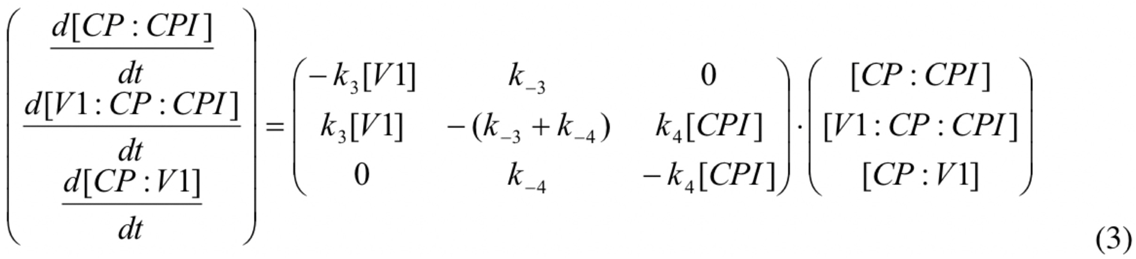

where ***CP*** is the vector of concentrations, 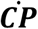 is the vector of time derivatives and ***M*** is the matrix of coefficients. For this system, two nonzero eigenvalues (*λ*_1_, *λ*_2_) can be obtained as the roots of the quadratic equation (Eq. 4) obtained from the condition |**M**-*λ***I**|=0, where **I** is the identity matrix (*25*),

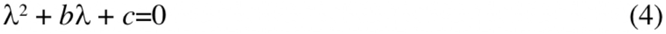

and where *b= k_4_ [CPI] - (k_-3_+k_-4_) + k_3_ [V-1]* and *c= (k_4_k_-3_ + k_3_k_4_ [V-1]) [CPI] + k_3_k_-4_ [V-1]*. These two eigenvalues describe the relative relaxation rates r_1_ and r_2_ as a function of [V-1] and [CPI] as shown in Eqs. 5.

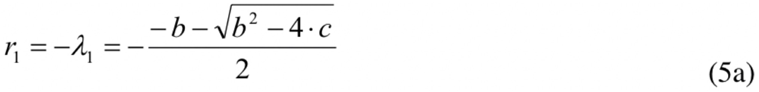

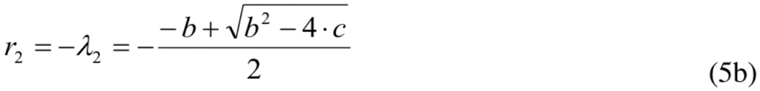

Eq. 5b was used to fit the dependencies of the observed rate on CPI-motif peptide concentration. Double-exponential fits of the stopped-flow data were used to obtain the rates for the larger-amplitude faster phase at each concentration of CPI-motif peptide, as described above. We noted that Eq. 5b contains four parameters (k_3_, k_-3_, k_4_ and k_-4_) that are highly correlated and cannot be obtained by direct fitting of the data. However, we reduced the number of fitting parameters to two (k_-3_ and k_-4_) by introducing values for K_d3_ and K_d4_ that were determined in independent equilibrium titration experiments (see Tables 1 and 2), and we defined bimolecular association constants in Eq. 5b as k_3_ = k_-3_/K_d3_ and k_4_ = k_-4_/K_d4_, respectively. Fitting was performed using MicroMath Scientist Software.

**Table 1.** Values for binding constants from experiments measuring the interaction of CPI-motif peptides with CP. Results from assays by fluorescence anisotropy titration (as in Figure 3) and by isothermal calorimetry (as in Figure 4). Corresponds to Reaction 1 in Figure 11. Fluorescence anisotropy values and errors are derived from global fits to combined data from two independent experiments; ITC values are from independent experiments, with errors from fitting. Fluorescence anisotropy experiments were also performed with unlabeled peptides, in competition with labeled ones, for three peptides. Those values, from single experiments, were as follows: CARMIL3, 1.4 ± 0.3 nM; CARMIL1, 20.6 ± 1.6 nM; Twf-1, 295 ± 45 nM. N.D. - not determined, because CapZIP peptide displayed aberrant fluorescence, preventing analysis.

|  | Fluorescence Anisotropy | Isothermal Calorimetry (ITC) |  |  |  |  |
| --- | --- | --- | --- | --- | --- | --- |
| CPI-Motif Peptide | $K_d$ obs (nM) | $K_d$ obs (nM) | $\Delta G^0_{obs}$ (kcal/mol) | $\Delta H_{obs}$ (kcal/mol) | $T\Delta S^0_{obs}$ (kcal/mol) | N |
| CARMIL3 | $10.0 \pm 2.1$ | $6.5 \pm 0.6$ | $-11.17 \pm 0.06$ | $-20.8 \pm 0.1$ | -9.7 | $0.927 \pm 0.003$ |
| | | $6.0 \pm 0.9$ | $-11.22 \pm 0.10$ | $-19.7 \pm 0.2$ | -8.5 | $1.030 \pm 0.005$ |
| WASHCAP | $8.5 \pm 0.8$ | $18 \pm 1$ | $-10.56 \pm 0.04$ | $-24.6 \pm 0.1$ | -14.1 | $0.985 \pm 0.004$ |
| | | $17 \pm 3$ | $-10.60 \pm 0.10$ | $-28.9 \pm 0.4$ | -18.3 | $0.971 \pm 0.009$ |
| CARMIL1 | $10.4 \pm 1.4$ | $27 \pm 2$ | $-10.32 \pm 0.05$ | $-16.4 \pm 0.2$ | -6.1 | $0.946 \pm 0.006$ |
| | | $15 \pm 1$ | $-10.66 \pm 0.05$ | $-14.7 \pm 0.1$ | -4.1 | $0.960 \pm 0.004$ |
| CapZIP | N.D. | $11 \pm 1$ | $-10.84 \pm 0.07$ | $-25.2 \pm 0.2$ | -14.4 | $1.010 \pm 0.005$ |
| | | $11 \pm 2$ | $-10.87 \pm 0.09$ | $-21.9 \pm 0.2$ | -11.0 | $1.060 \pm 0.007$ |
| CD2AP | $43 \pm 3$ | $23 \pm 2$ | $-10.41 \pm 0.04$ | $-17.4 \pm 0.1$ | -7.0 | $0.993 \pm 0.004$ |
| | | $19 \pm 2$ | $-10.54 \pm 0.07$ | $-21.5 \pm 0.2$ | -10.9 | $0.962 \pm 0.006$ |
| CIN85 | $59 \pm 2$ | $32 \pm 2$ | $-10.22 \pm 0.04$ | $-21.7 \pm 0.1$ | -11.5 | $1.010 \pm 0.005$ |
| | | $16 \pm 1$ | $-10.65 \pm 0.04$ | $-22.3 \pm 0.1$ | -11.6 | $1.000 \pm 0.004$ |
| | | $12 \pm 1$ | $-10.80 \pm 0.03$ | $-22.8 \pm 0.1$ | -11.9 | $0.989 \pm 0.002$ |
| CKIP-1 | $91 \pm 6$ | $100 \pm 6$ | $-9.55 \pm 0.04$ | $-16.4 \pm 0.2$ | -6.9 | $0.996 \pm 0.008$ |
| | | $36 \pm 3$ | $-10.15 \pm 0.05$ | $-14.0 \pm 0.1$ | -3.8 | $1.070 \pm 0.007$ |
| | | $29 \pm 2$ | $-10.28 \pm 0.04$ | $-15.7 \pm 0.1$ | -5.4 | $1.060 \pm 0.004$ |
| Twf-1 | $340 \pm 16$ | $295 \pm 13$ | $-8.91 \pm 0.03$ | $-10.2 \pm 0.1$ | -1.3 | $0.997 \pm 0.004$ |
| | | $310 \pm 12$ | $-8.88 \pm 0.02$ | $-10.2 \pm 0.1$ | -1.3 | $0.946 \pm 0.004$ |

**Table 2.** Values for binding and rate constants from experiments measuring the interactions of CP with V-1 and with F-actin. Values are means, and errors are standard deviations. Reactions 2 through 7 are illustrated in Figure 11. Values are derived from experiments similar to those in Figures 5 through 8. For Reaction 4, WASHCAP and CapZIP were not determined (N.D.) due to aberrant fluorescence of the CPI-motif peptides in the titration. For Reaction 5, CapZIP and Twf-1 were not determined (N.D.) due to effects of the CPI-motif peptides on actin polymerization in the absence of CP. The mutant peptide CARMIL1 / K987A R989A serves as a negative control in that these two amino-acid changes lead to a nearly complete loss of interaction of a CARMIL1 fragment with CP in actin polymerization assays (*17*). *This value is an approximation from the highest concentration of peptide tested; the kinetic model could not be used because the Kd for Reaction 4 was not determined due to the weak binding affinity of the mutant CARMIL1 peptide.

|  | **CP:V-1 with saturating CPI**<br>**(Reactions 2, 3)** |  | **CP:CPI with**<br>**saturating V-1**<br>**(Reaction 4)** | **CP:F-actin with**<br>**saturating CPI**<br>**(Reactions 5, 6)** | **CP:CPI, with**<br>**CP bound to**<br>**F-actin**<br>**(Reaction 7)** |
| --- | --- | --- | --- | --- | --- |
| **CPI-Motif Peptide** | **$K_d$ obs (nM)** | **$k_{off}$ obs ($s^{-1}$)** | **$K_d$ obs ($\mu M$)** | **$K_{CAP}$ obs (nM)** | **$K_d$ (nM)** |
| None | $21 \pm 3$ | $1.2 \pm 0.1$ | | $0.21 \pm 0.01$ | |
| CARMIL3 | $2100 \pm 300$ | $264 \pm 7$ | $0.63 \pm 0.05$ | $1.7 \pm 0.1$ | $80 \pm 17$ |
| WASHCAP | $3200 \pm 300$ | $209 \pm 4$ | N.D. | $2.7 \pm 0.1$ | $110 \pm 10$ |
| CARMIL1 | $1800 \pm 300$ | $211 \pm 6$ | $0.79 \pm 0.07$ | $1.3 \pm 0.1$ | $61 \pm 8$ |
| CapZIP | $3500 \pm 300$ | $31 \pm 3$ | N.D. | N. D. | |
| CD2AP | $3100 \pm 400$ | $33 \pm 7$ | $1.1 \pm 0.1$ | $2.9 \pm 0.1$ | $590 \pm 40$ |
| CIN85 | $1500 \pm 100$ | $150 \pm 5$ | $1.7 \pm 0.1$ | $0.93 \pm 0.02$ | $260 \pm 10$ |
| CKIP-1 | $1400 \pm 200$ | $55 \pm 4$ | $2.1 \pm 0.1$ | $3.1 \pm 0.1$ | $1300 \pm 100$ |
| Twf-1 | $93 \pm 35$ | $0.7 \pm 0.004$ | $0.78 \pm 0.05$ | N.D. | |
| CARMIL1/K987A R989A | $56 \pm 4$ | $1.36 \pm 0.01^*$ | N.D. | N.D. | |
| Method | fluorescence titration | stopped-flow kinetics | fluorescence anisotropy titration | actin capping polymerization | calculated |

### Actin Polymerization Assays

Pyrene labeling of actin was performed as described (*22*). F-actin seeds were prepared by adding 1 mM MgCl_2_, 1 mM EGTA, 50 mM KCl, and 2.1 *µ*M phalloidin to 2.1 *µ*M G-actin (5% pyrene label) and incubating at 25**°**C overnight (*26*). 0.5 *µ*M pyrene-labeled F-actin seeds, in the presence of 0, 0.75, 2, 5, 10, 25, and 50 nM CP, with and without 20 *µ*M CPI-motif peptides, were incubated at 25**°**C for 30 min. Based on the binding affinities of the CPI-motif peptides for CP, a single-site binding model indicates that at concentrations of CP in the assay, the CPI-motif peptides are saturated at 20 *µ*M. WASHCAP CPI peptide was tested at 1 *µ*M because at higher concentrations the peptide interfered with polymerization. To initiate polymerization, 1 mM MgCl_2_ and 1 mM EGTA were added to pyrene-labeled G-actin; the Mg^2+^-primed actin was then added at 1.5 *µ*M to the F-actin seed mixture. Elongation rates were measured using time-based scans on a plate reader (Biotek Synergy H4 Hybrid Multi-Mode Microplate Reader with Gen5 software, BioTek Instruments, Winooski, VT) at 25°C with excitation at 365 nm and emission detected at 407 nm.

The apparent K_d_ of CP binding to the barbed end (K_CAP_) was calculated from the rate constants for CP binding to the barbed end. The rate constants were obtained from experiments in triplicate, which were fit to a model based on earlier work describing the kinetics of actin polymerization ((*27, 28*) using KinTek Explorer Software V8.0 (KinTek Corporation, Snow Shoe, PA). The nine reactions in the model are shown below, with A = actin monomer, A2 = actin dimer, A3 = actin trimer, A4 = actin tetramer, A5 = actin pentamer, Nb = barbed end, Np = pointed end, CP = capping protein:

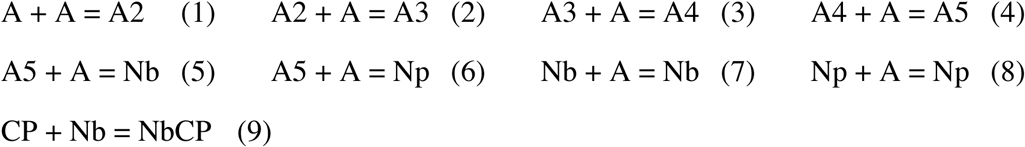

Rate constants for actin elongation, obtained from (*29*), were as follows: k_+5_ = 1.16 × 10^-5^ M^-1^s^-1^, k_+7_ =1.16 × 10^-5^ M^-1^s^-1^, k_-7_ = 1.4 s^-1^. Other rate constants were determined from fitting results of polymerization experiments in the absence of CP, CPI-motif peptides, and F-actin seeds. We assumed that reactions 5 and 6 could only proceed in the forward direction, with reverse reaction rates constrained to zero, and we assumed that the association reaction in step 9 was diffusion-limited at 10^9^ M^-1^s^-1^. Values were as follows: k_+1_ = 9.47 × 10^-7^ M^-1^s^-1^, k_-1_ = 1.24 s^-1^, k_+2_ = 1.01 × 10^-9^ M^-1^s^-1^, k_-2_ = 1.25 s^-1^, k_+3_ = 7.27 × 10^-8^ M^-1^s^-1^, k_-3_ = 1.33 s^-1^, k_+4_ = 5.72 × 10^-8^ M^-1^s^-1^, k_-4_ = 1.31 s^-1^, k_-5_ = 0 s^-1^, k_+6_ = 9.04 × 10^-7^ M^-1^s^-1^, k_-6_ = 0 s^-1^, k_+8_ = 9.08 × 10^-7^ M^-1^s^-1^, k_-8_ = 0.804 s^-1^, k_+9_ = 1.0 × 10^9^ M^-1^s^-1^.

To fit experimental data and determine K_CAP_, the rate constants above were kept constant, and Np and Nb were constrained to equal values determined from polymerization experiments without CP or CPI-motif peptides. k_-9_ was solved by simultaneous fitting of the polymerization data for all concentrations of CP within an experiment. K_CAP_ was calculated by dividing k_-9_ by k_+9_. When high concentrations of CP (>10 nM) caused filament nucleation from actin subunits alone, those results were omitted from the global fit.

## RESULTS

### Conservation of CPI-Motif Sequences

We asked whether CPI-motif sequences in different proteins are conserved across evolution, which might suggest functional differences in cells. We selected CPI-motif regions from vertebrate proteins currently known to contain CPI motifs, including CARMIL, CD2AP, CIN85, CKIP, CapZIP, WASHCAP, and Twinfilin; and we performed multiple sequence alignment with ClustalW. The alignment is shown in Figure 1A, with expanded views in Supplemental Figure 1, panels a to k. Conservation of sequence within families and the relationships among families are illustrated in an unrooted phylogenetic tree derived from the alignment (Figure 1B). Organism names are displayed on a separate unrooted phylogenetic tree with a horizontal orientation (Supplemental Figure 2).

**Figure 1.**
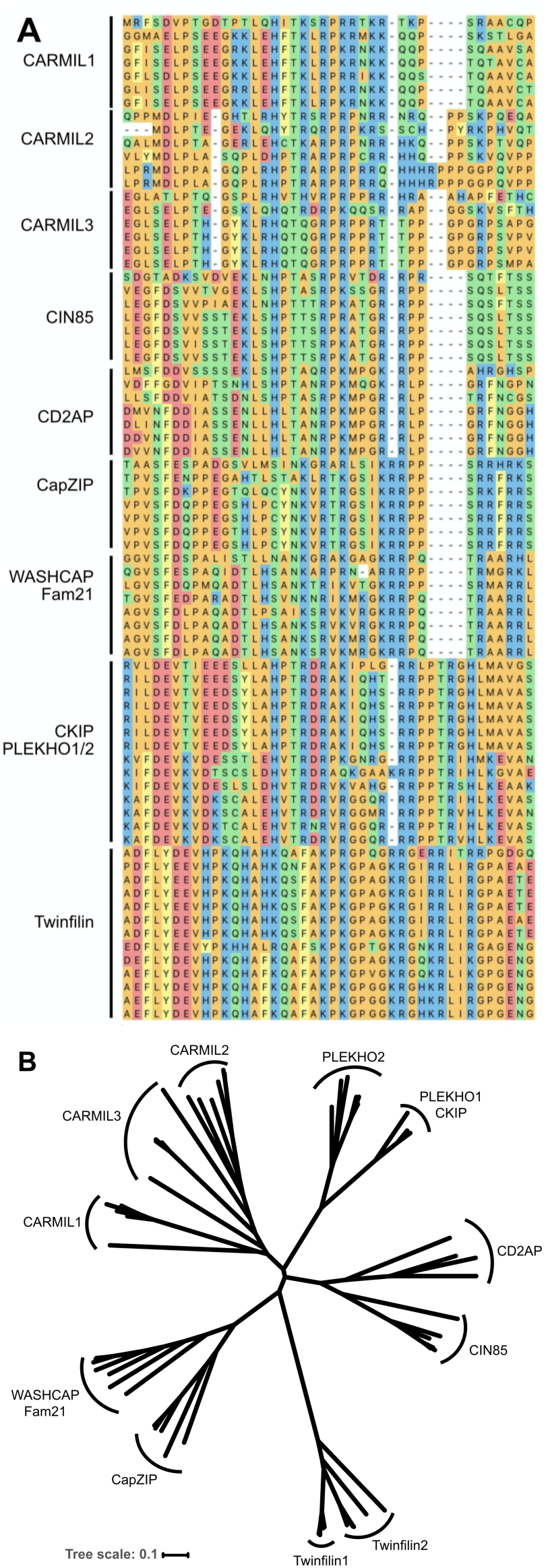
Phylogenetic analysis of CPI-motif peptides. (A) Multiple sequence alignment of CPI-motif peptides. ClustalW sequence alignment of CPI-motif peptides from representative vertebrates. Color scheme by chemistry, default in DNASTAR: yellow - aromatic, red - acidic, blue - basic, orange - nonpolar, green - polar. Detailed views and sequence logos are presented in Supporting Information as Supplemental Figure 1 panels A to K. Peptides were derived from the CPI-motif protein families with the following accession numbers: CARMIL1: Zebrafish ALI93829, Chicken NP_001152842, Koala XP_020842531, Camel XP_010978439, Mouse NP_081101, Human Q5VZK9; CARMIL2: Zebrafish ALI93830, Frog XP_004913561, Chicken XP_015134656, Koala XP_020844996, Mouse NP_001344262, Human NP_001013860; CARMIL3: Zebrafish ALI93831, Frog NP_001121429, Koala XP_020822585, Camel XP_010994108, Mouse NP_001019816, Human Q8ND23; CIN85: Zebrafish XP_021326013, Frog XP_002935234, Chicken XP_015157987, Koala XP_020845587, Camel XP_010985735, Mouse NP_001129199, Human Q96B97; CD2AP: Zebrafish NP_001008583, Frog NP_001086432, Chicken NP_001305332, Koala XP_020849556, Camel XP_010993738, Mouse NP_033977, Human NP_036252; CapZIP: Zebrafish NP_001038887, Frog BAR45528, Chicken NP_001025960, Koala XP_020830347, Camel EPY73831, Mouse NP_848708, Human NP_443094; WASHCAP: Zebrafish subunit 2C XP_005156762, Frog subunit 2C XP_017951133, Chicken subunit 2C NP_001012611, Koala subunit 2C XP_020850229, Camel subunit FAM21-like XP_010983602, Mouse subunit 2C XP_006506277, Human subunit 2C NP_001162577; Human subunit 2A NP_001005751; CKIP-1: Zebrafish BAF62166, Frog XP_002938507, Chicken NP_001026527, Koala XP_020860003, Camel XP_010995007, Mouse NP_075809, Human NP_057358; CKIP-2: Zebrafish XP_002662798, Frog NP_001106552, Chicken XP_025009763, Koala XP_020839263, Camel XP_010991846, Mouse NP_694759, Human NP_079477; Twinfilin-1: Zebrafish AAI53600, Frog AAH88597, Chicken NP_001265024, Koala XP_020829343, Camel XP_010987877, Mouse NP_032997, Human NP-002813; Twinfilin-2: Zebrafish NP_001018486, Frog NP_001123417, Chicken NP_001025760, Camel XP_014410799, Mouse NP_036006, Human NP_009215. (B) Phylogenetic analysis of CPI-motif peptide families. Unrooted phylogenetic tree of aligned CPI-motif peptides from Figure 1A. Sequences are conserved within a family, and families are distinct from one another. Organism names are presented in an unrooted version of the tree in Supporting Information as Supplemental Figure 2. The tree scale represents the number of differences between sequences, 0.1 corresponding to 10% differences between two sequences.

### Structures and Surface Contacts for Interactions of CP with F-Actin, V-1 and CPI-motif

The binding sites on CP for V-1 and F-actin are overlapping but not identical (*7, 30–32*). The binding site on CP for CPI motifs is quite distinct from the V-1 binding site and the F-actin binding site (*1, 7, 19*). To illustrate these points, structures of complexes are shown in Figure 2A, and contact surfaces are shown in Figure 2B. In Figure 2B, the actin contact surface is red, the V-1 surface is yellow, the overlap between the two is orange, and the binding site for CPI motifs is in blue.

**Figure 2.**
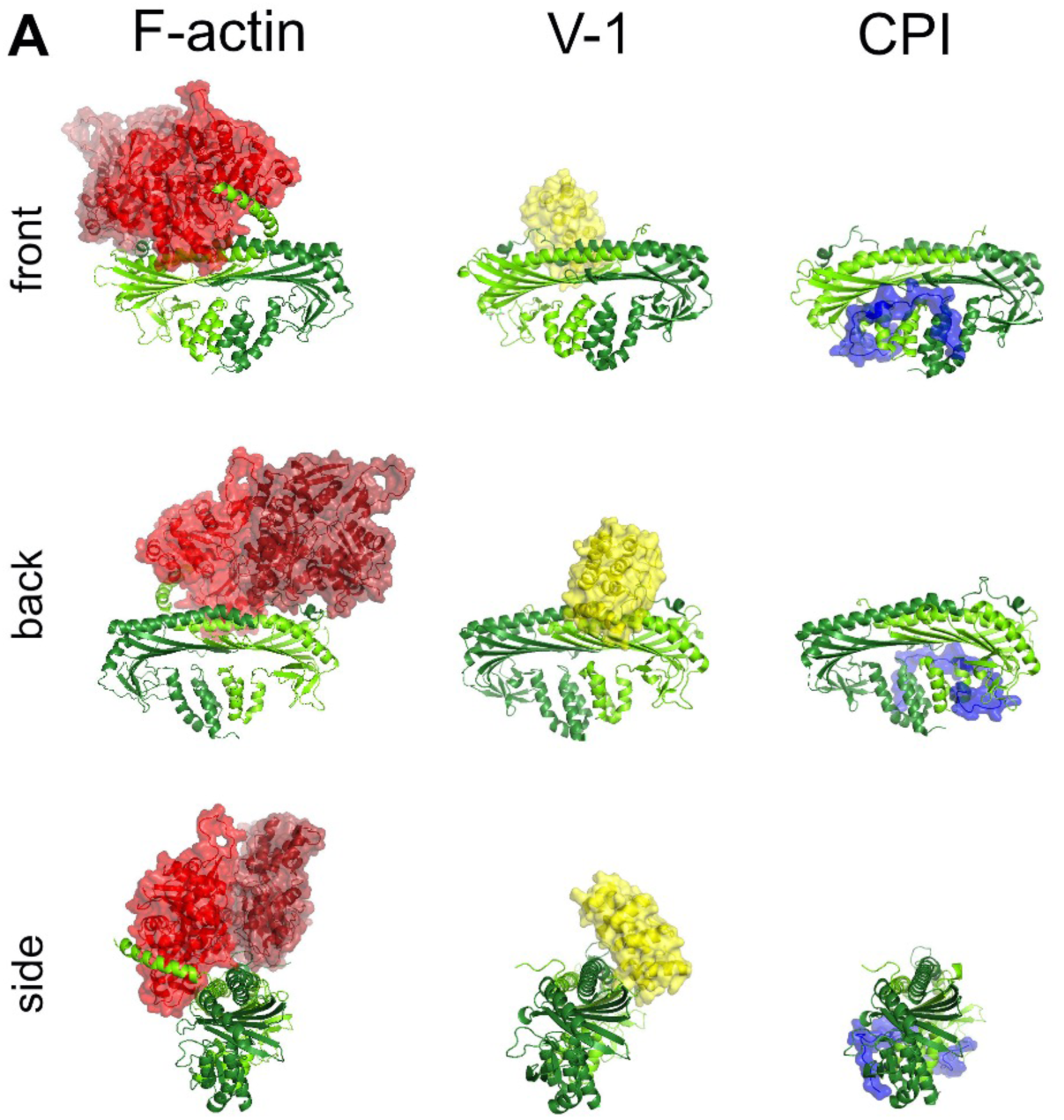
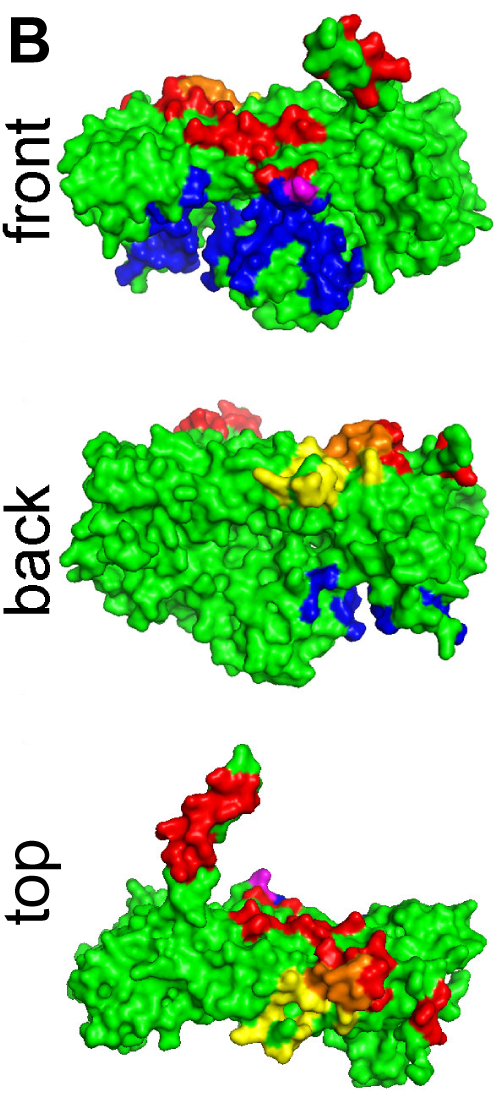
Structures of Capping Protein, Binding Partners and Contacts. (A) Structures of Capping Protein with Binding Partners. Capping protein (CP) bound to actin (red), V-1 (yellow) and CARMIL1 CPI region (blue). V-1 sterically inhibits CP binding to actin, while CPI-motifs allosterically alter CP binding to actin. The actin-CP structural model was prepared by Dr. Roberto Dominguez (University of Pennsylvania). The model combines the structure of CPα1β2 from a cryo-EM structure of dynactin (PDB ID: 5ADX; Urnavicius et al., Science 2015) with structures of two conventional actin protomers (PDB ID: 6DJM; Chou and Pollard PNAS 2019) replacing the Arp1 subunits of dynactin. The CP:V-1 structure (PDB ID: 3 AAA; Takeda et al., PLoS Biol. 2010) and the CP:CPI structure (PDB ID: 3LK3; Hernandez-Valladares et al., Nat. Struct. Mol. Biol. 2010) were generated from published X-ray crystal structures prepared using CPα1β1. (B) Contact Surfaces for Capping Protein Binding Partners. CP and binding partner surface contacts were analyzed using the structures in Figure 2A. Contact residues in CP within 3.9 Å of the ligand are indicated here. Contact surfaces for F-actin (red), V-1 (yellow) and CARMIL 1 CPI (blue) are shown. Overlapping contact residues on CP for actin and V-1 are shown in orange. Several of the overlapping residues are important for actin binding, demonstrated by biochemical assays and structural models for CP binding to the barbed end of an actin filament (*31, 33*). A single overlapping contact residue in CP for actin and CARMIL 1 CPI is shown in purple, while the majority of the contact surface for CPI binding is distal to the actin binding site.

In light of the conserved differences among the CPI-motif sequences of the families and the differences between the binding sites for F-actin and V-1, we compared the CPI-motif families in measurements of binding affinity for CP and allosteric effects on the interactions of V-1 and F-actin with CP.

### Binding of CPI-motif Peptides to CP

To determine the binding affinity of CPI-motif peptides for CP, we titrated CP into fluorescently labeled CPI-motif peptides. We monitored fluorescence anisotropy because little or no changes in fluorescence intensity were observed. The data were fit to a single-site binding model. Figure 3 shows titrations and fits for all CPI-motif peptides tested. K_d_ values for these and other CPI-motif peptides are listed in Table 1. For three CPI-motif peptides, CARMIL3, CARMIL1 and Twinfilin, we confirmed the result by measuring the binding affinity with unlabeled peptides, using a competition approach (Table 1 Legend).

**Figure 3.**
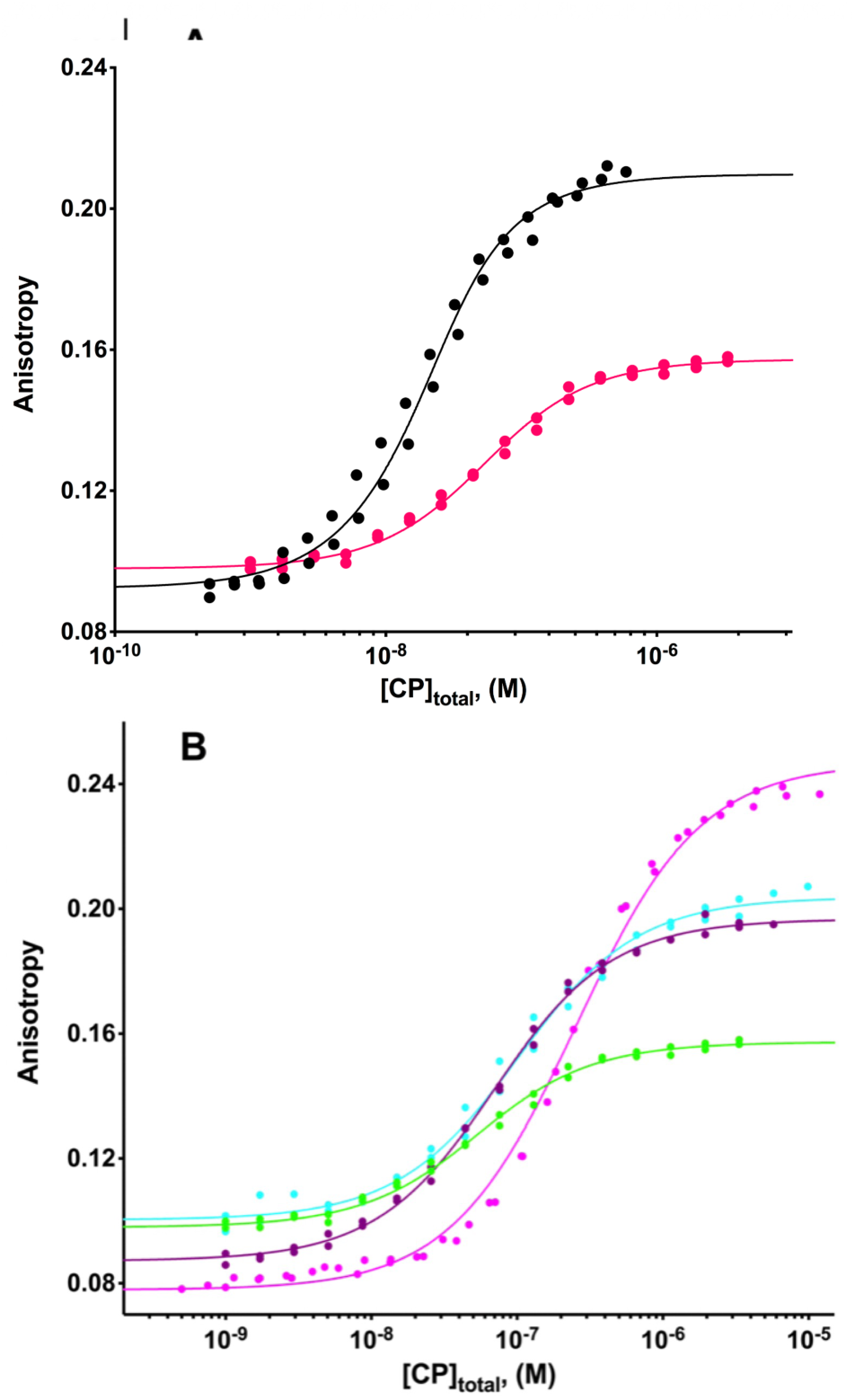
Binding affinity of CPI-motif peptides for CP by fluorescence anisotropy. Anisotropy of TAMRA-labeled CPI-motif peptides plotted versus total concentration of CP. (A) WASHCAP, black; CARMIL3, blue; CARMIL1, red. (B) CD2AP, green; CIN85, purple; CKIP-1, cyan; Twf-1, magenta. The points represent data from experiments in duplicate, and the lines are the best fits to a single-site binding model. The fitted values of the K_d_ for CPI-motif peptides binding to CP are listed in Table 1.

The fluorescence anisotropy titrations were well-behaved and consistent with a single-site binding model. Conditions were 100 mM KCl and 20 mM MOPS pH 7.2. We also measured binding over a range of pH and salt concentrations for two CPI-motif peptides, CARMIL1 and CARMIL3, as shown in Supplemental Figure 3. For both peptides, binding affinity decreased with decreasing pH and higher salt concentration. Competition experiments with unlabeled peptide were also performed for CARMIL1 at different pH and salt conditions, as described in the legend to Supplemental Figure 3.

We also measured the binding of CPI-motif peptides to CP by isothermal titration calorimetry (ITC), titrating unlabeled CPI-motif peptides into CP. Representative titrations and fits for two CPI-motif peptides, from CARMIL1 and CD2AP, are shown in Figure 4. Values for K_d_, ΔG^0^, ΔH, and TΔS^0^ for all CPI-motif peptides examined are listed in Table 1. The values for the binding constants from fluorescence anisotropy and ITC were similar for all CPI-motif peptides, differing by no more than 3-fold (comparing global-fit fluorescence anisotropy values with means of ITC values). The ITC results revealed the binding reaction to be driven by enthalpy, with a stoichiometry of 1:1; these conclusions are consistent with the chemical nature of the binding interface as revealed by structural studies of the complexes (*1, 7, 19*).

**Figure 4.**
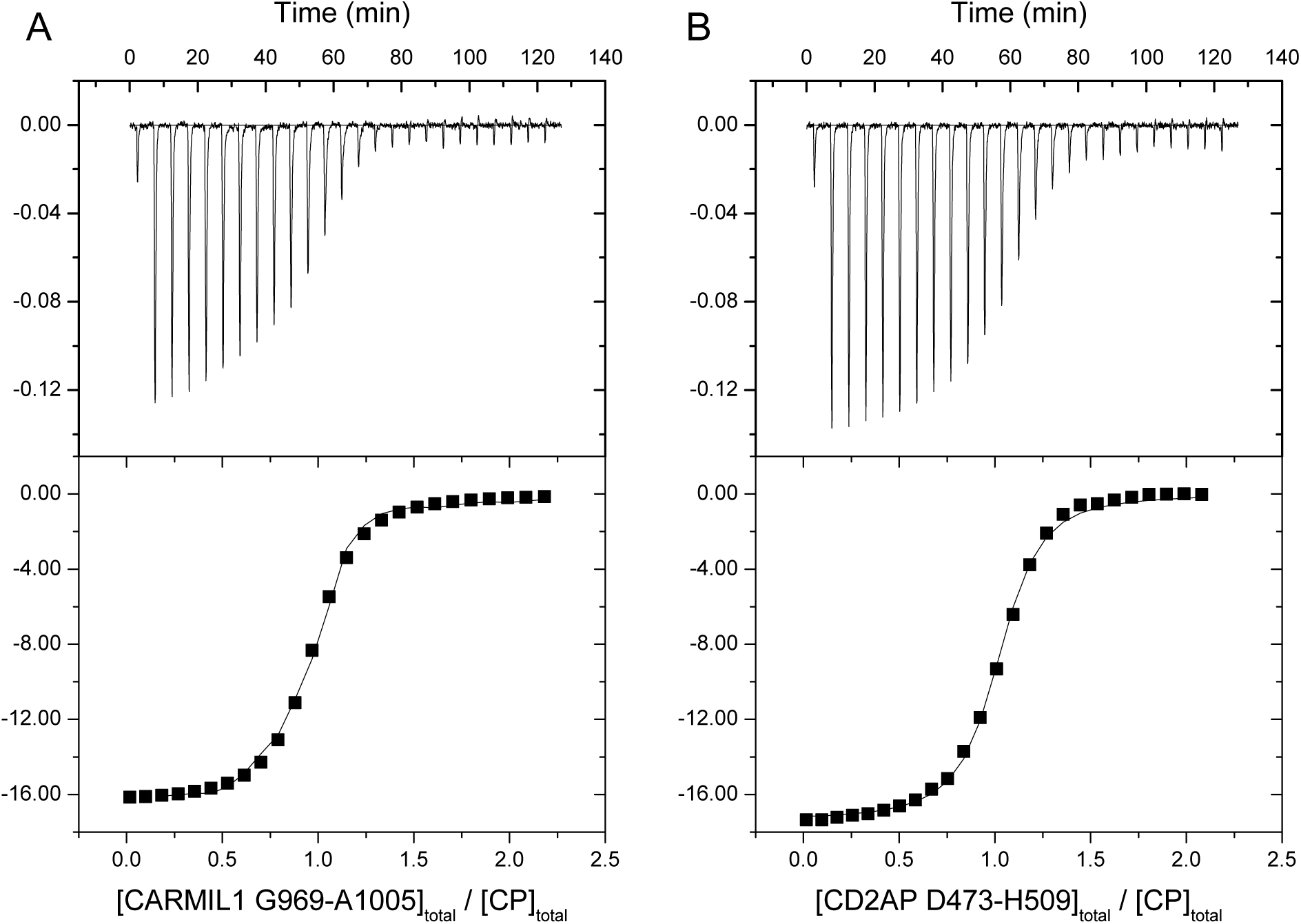
ITC of CPI-motif peptides binding to CP. Two representative examples are shown. Upper panels: Titration of CPI-motif peptides into CP with raw data plotted as the heat signal versus time. Lower panels: Smooth curve shows the best fit of the data to a single-site binding model. (A) Titration of CARMIL1 into CP. N = 0.946 ± 0.006, K_d_ = 27 ± 1 nM, ΔH^0^ = −16.4 ± 0.2 kcal/mol, TΔS^0^ = −6.1 kcal/mol. (B) Titration of CD2AP into CP. N = 0.993 ± 0.004, K_d_ = 23 ± 2 nM, ΔH^0^ = −17.4 ± 0.1 kcal/mol, TΔS^0^ = −6.9 kcal/mol.

### Effect of V-1 on Binding of CPI-motif Peptides to CP

In cells, a substantial fraction of CP is likely to exist in 1:1 complex with V-1, freely diffusing about the cytoplasm (*8*). Therefore, a molecule of CP that encounters a CPI motif is likely to be bound to a molecule of V-1, thus forming a ternary complex. The binding of V-1 and CPI motifs to CP are mutually antagonistic, in an allosteric mechanism (*8*). Thus, we investigated the interaction of CPI-motif peptides with the CP:V-1 complex.

We monitored the increase in fluorescence anisotropy as CP was titrated into a solution of fluorescently labeled CPI-motif peptide, in the presence of saturating concentrations of V-1. The data were fit to a single-site binding model. Representative titration curves are shown in Figure 5, for CARMIL1 and CD2AP. The fitted K_d_ value for CARMIL1 was 790 ± 70 nM; in the absence of V-1, the corresponding K_d_ was 10.4 ± 1.4 nM (Table 1). For CD2AP, the K_d_ was 1.1 ± 0.1 *µ*M; in the absence of V-1 the value was 43 ± 3 nM (Table 1). The values for all the CPI motifs examined are listed in Table 2, under Reaction 4. For every CPI motif, the presence of V-1 decreased the binding affinity of the CPI motif for CP. The magnitude of this effect on K_d_, attributable to V-1, varied from 100-fold to 500-fold (Table 2, under Reaction 4). The implications of these observations are discussed below.

**Figure 5.**
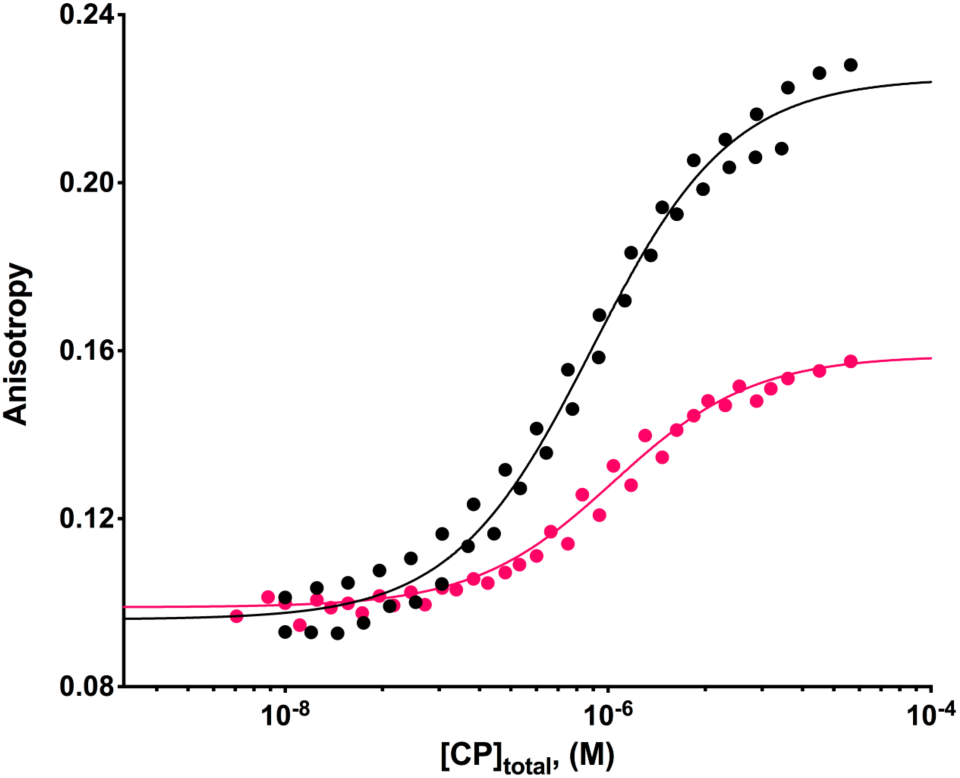
Effect of V-1 on the binding affinity of CPI-motif peptides for CP by fluorescence anisotropy. Two representative examples are shown. Anisotropy of TAMRA-labeled CARMIL1 in the presence of 20 *µ*M and 50 *µ*M V-1 (black) and CD2AP in the presence of 50 *µ*M V-1 (red) is plotted versus total concentration of CP. The data points are from individual duplicate experiments, and the lines are the best fits to a single-site binding model. Fitted values of the K_d_ for CPI-motif peptides binding to CP in the presence of V-1 are listed in Table 2.

### Effect of CPI-motif Peptides on V-1 Binding to CP

One cellular pool of CP with potentially great physiological significance is that found as a 1:1 complex with a CPI-motif protein that has been targeted to a membrane (*11, 14*). Cell studies suggest that this pool of CP is part of the active fraction of CP, available to cap barbed ends of actin filaments that are near membranes (*15, 34*). These barbed ends may have arrived at the membrane by growth of barbed ends that were nucleated by Arp2/3 complex or formins, whose activity can depend on membrane-associated signals and regulators. This pool of CP may thus attach barbed ends to membranes and stabilize those barbed ends against actin subunit addition and loss.

Another potential outcome for this pool of CP, which is bound to a CPI-motif protein at a membrane, could be for V-1, which is freely diffusible and soluble, to bind to CP and thus sterically block its binding site for actin. To investigate this possibility, we examined the effect of CPI motifs on the binding of V-1 to CP.

We measured the effect of CPI-motif peptides on the binding affinity of V-1 for CP by titrating CP into fluorescently labeled V-1, in the presence and absence of saturating concentrations of CPI-motif peptides. Fluorescence intensity increased when fluorescent V-1 bound to CP, so we monitored intensity instead of anisotropy. The data from duplicate experiments were fit to a single-site binding model. Representative titrations, for CARMIL1 and CD2AP, are shown in Figure 6. The fitted K_d_ values for V-1 binding to CP were as follows: 21 ± 2 nM in the absence of CPI peptides, 1.8 ± 0.3 *µ*M in the presence of saturating CARMIL1, 56 ± 4 nM in the presence of saturating CP-binding mutant CARMIL1 (G969-A1005 / K987A R989A) (*1*) and 3.1 ± 0.4 *µ*M in the presence of saturating CD2AP. The presence of mutant CARMIL1 (G969-A1005 / K987A R989A) had a very small effect on V-1 binding, as expected. V-1 binding constants for all the CPI-motif peptides examined are listed in Table 2, under Reaction 3. We also measured the binding of V-1 to CP in the absence of CPI-motif peptides by ITC, titrating V-1 into CP. The binding affinity was similar to that determined by fluorescence titration binding experiments, with a K_d_ of 13.7 ± 1.8 nM. The binding is driven by enthalpic and entropic terms (ΔH^0^ = −3.9 kcal/mol; TΔS^0^ = 6.8 kcal/mol; N = 1.030 ± 0.005), in contrast to the CPI-motif peptide binding reaction for which the enthalpy term outweighs the opposing entropy term.

**Figure 6.**
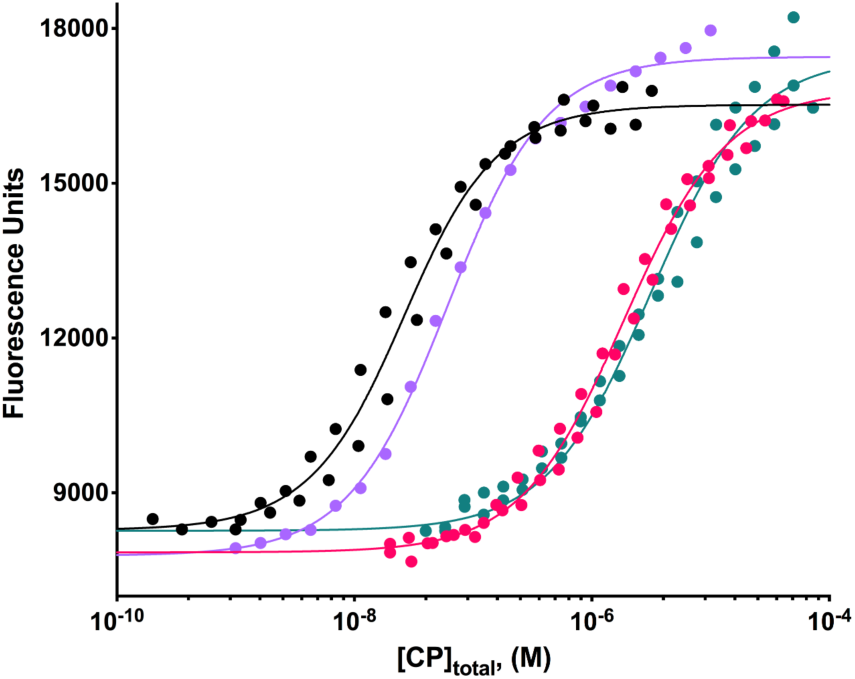
Effect of CPI-motif peptides on the binding affinity of V-1 for CP by fluorescence spectroscopy. Representative examples are shown. The fluorescence intensity of TAMRA-labeled V-1 is plotted versus the total concentration of CP. Experiments performed in the absence of any CPI-motif peptide (black), or the presence of saturating concentrations of CARMIL1 (red), mutant CARMIL1 (K987A R989A) (lavender) or CD2AP (teal). The CPI-motif peptides were present at 50 *µ*M, at least 150-fold greater than the K_d_ for each CPI-motif-peptide binding to CP in the absence of V-1. The points represent data from experiments in duplicate, except for one replicate for CARMIL1 (K987A R989A). The lines are the best fits to a single-site binding model. The fitted values of the K_d_ for V-1 binding to CP in the presence of CPI-motif peptides are listed in Table 2.

### Effect of CPI-motif Peptides on Dissociation of V-1 from CP

Actin assembly in a living cell can appear to be at steady state, but the system is in fact far from equilibrium. The actin cytoskeleton consumes large quantities of ATP in processes that drive the dynamic assembly and disassembly of actin filaments, and these dynamics contribute to movements in cells. Many aspects of actin assembly dynamics are proposed to involve the capping or uncapping of barbed ends of actin filaments, and actin assembly dynamics operate on the time scale of seconds; therefore, the time scale of the regulation of CP is important. Most of the CP in the cytoplasm is unable to interact with F-actin due to its association with V-1; however, CPI-motif proteins promote dissociation of the CP:V-1 complex, and this liberates V-1 from CP, thus activating the capping activity of CP. We therefore investigated the kinetics of V-1 dissociation from the CP:V-1 complex induced by different CPI-motif-protein peptides. In previous reports, the CPI-motif regions of CARMIL1 and twinfilin were found to increase the dissociation rate of V-1 from CP (*7, 8, 21*).

We compared the abilities of CPI-motif peptides from different protein families to induce dissociation of V-1, using a stopped-flow fluorescence assay. TAMRA-V-1 was allowed to form a complex with CP, and then mixed at time zero with an excess concentration of unlabeled V-1. The resulting decrease in fluorescence reflected dissociation of TAMRA-V-1 from CP, as shown in Figure 7A. We titrated the reaction with increasing concentrations of CPI-motif peptides, by adding them to the solution of unlabeled V-1. The observed relaxation rate for V-1 dissociation increased with the concentration of CPI-motif peptide, as shown in Figure 7B. The CP binding mutant CARMIL1 (K987A R989A) had only a minimal effect on V-1 dissociation. The relaxation rate for V-1 in response to all the CPI-motif peptides examined are listed in Table 2 under Reaction 3 (stopped-flow kinetics).

**Figure 7.**
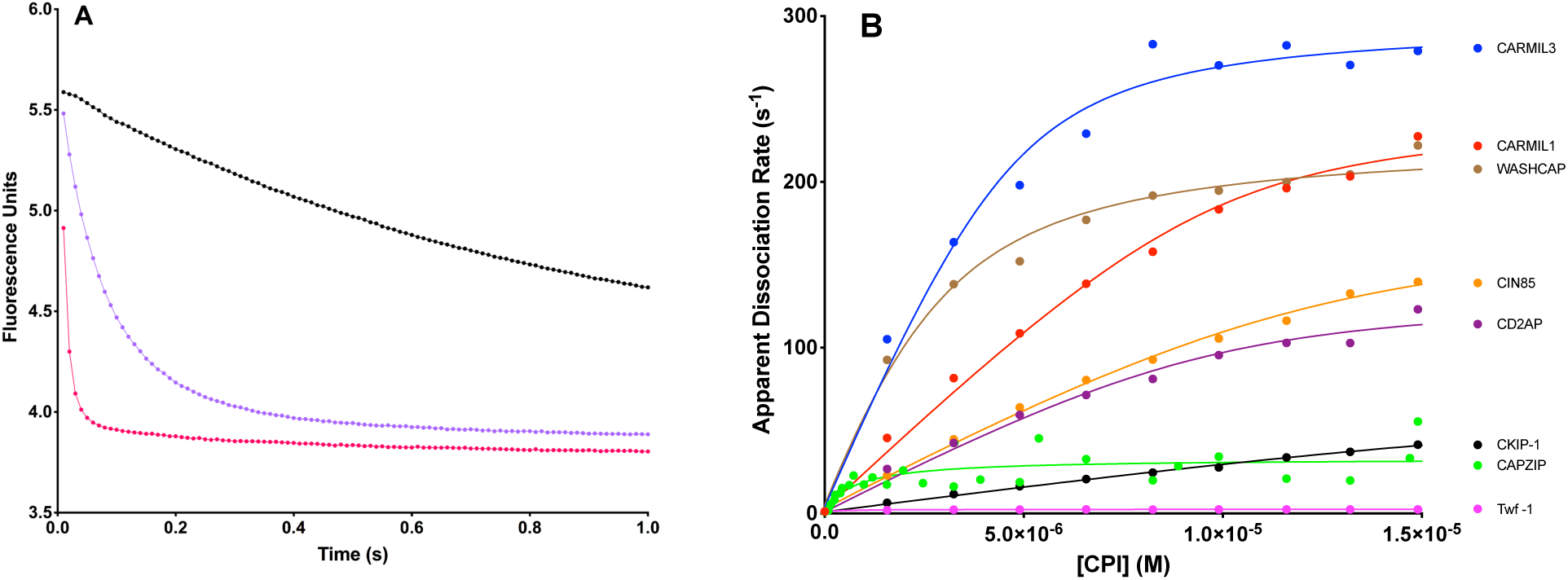
Effect of CPI-motif peptides on dissociation rate of V-1 from CP by stopped-flow fluorescence spectroscopy. (A) Fluorescence intensity plotted versus time, for fluorescently labeled V-1 dissociating from CP upon addition of unlabeled V-1 at time zero. Three representative examples are shown: control with only unlabeled V-1 (black), CARMIL1 peptide at 0.5 *µ*M (lavender) and 5 *µ*M (red). The points represent experimental data, and the lines are the best fits to a double-exponential decay model. The value of the smaller-amplitude step of the double exponential was 90- to 1700-fold less than the value of the larger-amplitude step, and it showed little dependence on the concentration of CPI motif. Thus, we report only the fitted values for the rate of the larger-amplitude process here. The apparent dissociation rates for V-1 from CP in these examples were as follows: 1.23 ± 0.03 s^-1^ in the absence of CPI-motif peptide, 13.6 ± 0.1 s^-1^ in the presence of 0.5 *µ*M CARMIL1, and 92 ± 1 s^-1^ in the presence of 5 *µ*M CARMIL1. (B) Apparent dissociation rates of V-1 from CP, obtained from experiments such as those in panel A, plotted versus the concentration of CPI-motif peptide. Shown are the following: CARMIL3, blue; CARMIL1, red; WASHCAP, brown; CIN85, orange; CD2AP, purple; CKIP-1, black; CapZIP, green; Twf-1, magenta. The points are values from experiments, and the solid lines are simulations of best fits to a kinetic model. V-1 dissociation rate constants in the absence and presence of CPI-motif peptides, are listed in Table 2, under Reactions 2 and 3.

### Effect of CPI-motif Peptides on CP Binding to Actin Barbed Ends (Capping)

To measure and compare the effects of different CPI-motif peptides on the affinity of CP for the barbed end, we performed actin polymerization experiments. We assayed the fluorescence intensity of pyrene-labeled actin over time, seeding the reaction with pre-formed barbed ends from actin filaments. We used a range of CP concentrations, in the absence and presence of saturating concentrations of CPI-motif peptides. The results from experiments in triplicate were fit to a kinetic model for actin polymerization. The model produced fitted values for rate constants of CP binding to the barbed end, allowing us to calculate K_d_s for the barbed end, K_CAP_. Reaction time courses with different concentrations of CP and no CPI-motif peptide are shown in Supplemental Figure 4. Representative curves for effects of CARMIL1 and CD2AP peptides on actin capping by CP, measured in this assay, are shown in Figure 8. Values for K_CAP_ in the absence and presence of all CPI-motif peptides examined are listed in Table 2 under Reactions 5 and 6. Values were not determined for CapZIP and Twf-1 because these CPI-motif peptides had effects on actin polymerization in the absence of CP.

**Figure 8.**
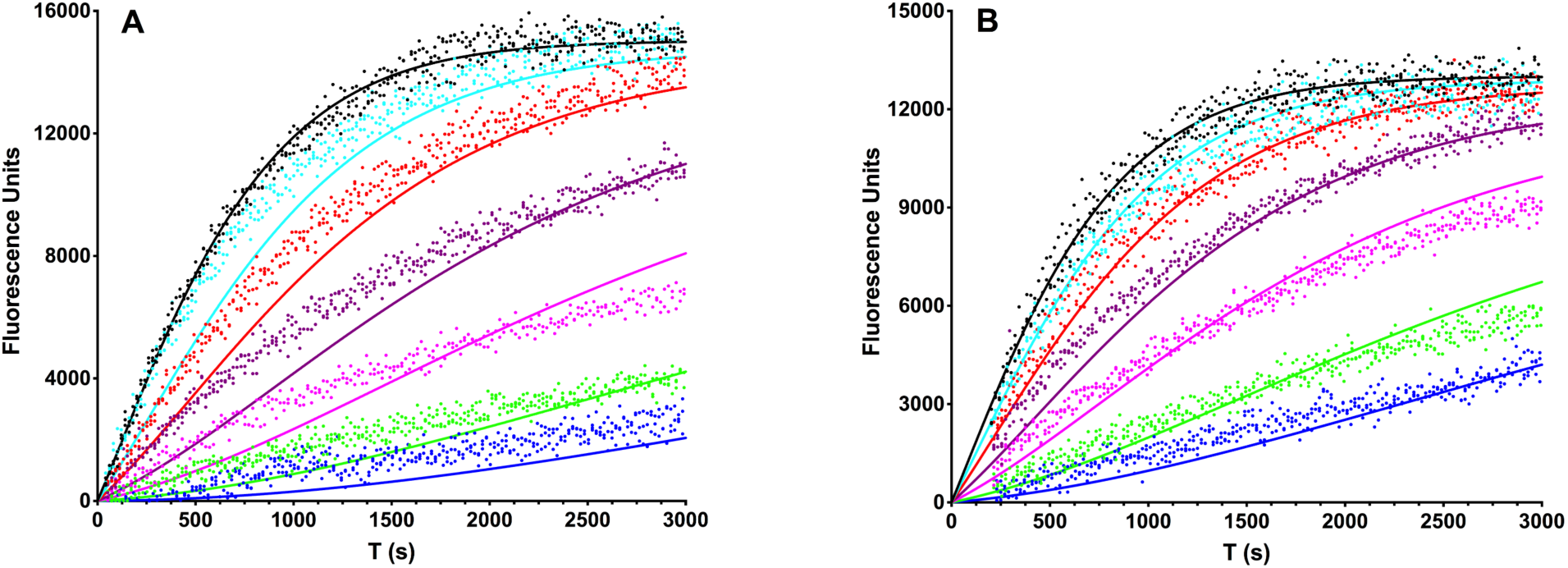
Effect of CPI-motif peptides on the affinity of CP for the barbed end by pyrene actin fluorescence spectroscopy. Pyrene-actin fluorescence is plotted versus time. The points represent data from experiments in triplicate, and the lines are the best simultaneous (global) fits to an actin polymerization model. (A) Experiments were performed in the presence of saturating (20 *µ*M) CARMIL1, and the following concentrations of CP: 0 (black), 0.75 (cyan), 2 (red), 5 (purple), 10 (magenta), 25 (green), and 50 nM (blue). The fitted value for the K_d_ of CP binding to the barbed end (K_CAP_) in the presence of CARMIL1 was 1.3 ± 0.1 nM. (B) Experiments as in panel A with saturating (20 *µ*M) CD2AP. K_CAP_ in the presence of CD2AP was 2.9 ± 0.1 nM.

### Comparison of Order of Effects

We hypothesized that the conservation of primary sequence across evolution, as found above and illustrated in Figure 1, reflects specific and distinct differences in one or more of the biochemical properties measured here, including actin binding and V-1 binding. If this were the case, we reasoned that the different CPI-motif peptides, would show a lack of correlation among the different assays. To address this question, we asked how well activity in one assay correlated with activity in another assay.

First, we compared the binding affinity of the CP:CPI complex for V-1 with the binding affinity of the CPI motif for CP (Figure 9A); this plot reveals only a rough inverse correlation among the K_d_ values. We also compared the rate of dissociation of V-1 from the CP:CPI complex with the binding affinity of the CPI motif for CP (Figure 9B); this plot also shows only a rough inverse correlation. These results reveal substantial differences among the CPI motifs, considering them alone and with respect to each other. The results indicate that greater energy of binding of CPI to CP is accompanied by greater loss of energy of binding of V-1 to CP.

**Figure 9.**
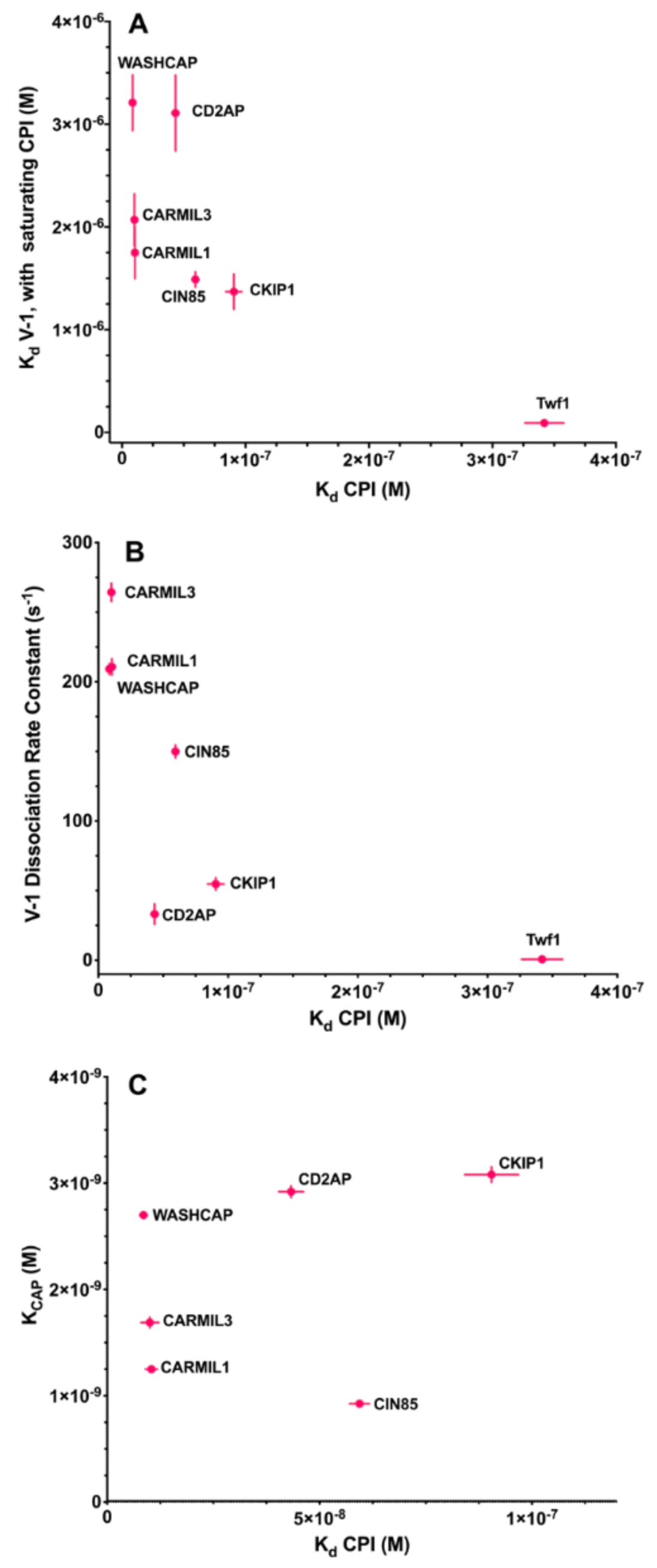
Comparison of CP:V-1 and CP:Actin interactions with CP:CPI binding affinity. Three plots comparing allosteric effects on the actin- and V-1-binding upper surface of CP (mushroom cap) with binding of a CPI-motif peptide to the lower portion (mushroom stalk) of CP. Plotted points are values listed in Tables 1 and 2. CPI motifs are labeled on the graphs. The error bars are derived from the experimental data; and when error bars are not visible, the error was less than the radius of the symbol. (A) Binding affinity of V-1 for CP:CPI (CP in presence of saturating CPI) plotted versus binding affinity of CP for CPI motif. (B) Dissociation rate constants of V-1 from CP:CPI:V-1 complex plotted versus binding affinity of CP for CPI motif. (C) Binding affinity of actin (F-actin barbed end) for CP plotted versus binding affinity of CP for CPI motif.

Next, we compared the binding affinity of the CP:CPI complex for F-actin, based on the apparent K_d_ for actin capping, with the binding of the CPI motif for CP (Figure 9C). The values show little or no correlation, providing another example of differences among the CPI motifs that are not proportional to the binding affinity of the CPI motif for CP in a straightforward relationship.

Finally, we compared the effects of CPI motif peptides on actin capping compared with their effects on V-1 interactions (Figure 10). The binding affinity of the CP:CPI complex for V-1 correlated relatively well with the affinity of CP:CPI complex for F-actin, based on the apparent K_d_ for actin capping, with the exception of CKIP-1 (Figure 10A). The rate of dissociation of V-1 from the CPI:CP:V-1 complex showed almost no correlation when compared with the affinity of the CP:CPI complex for F-actin (Figure 10B).

**Figure 10.**
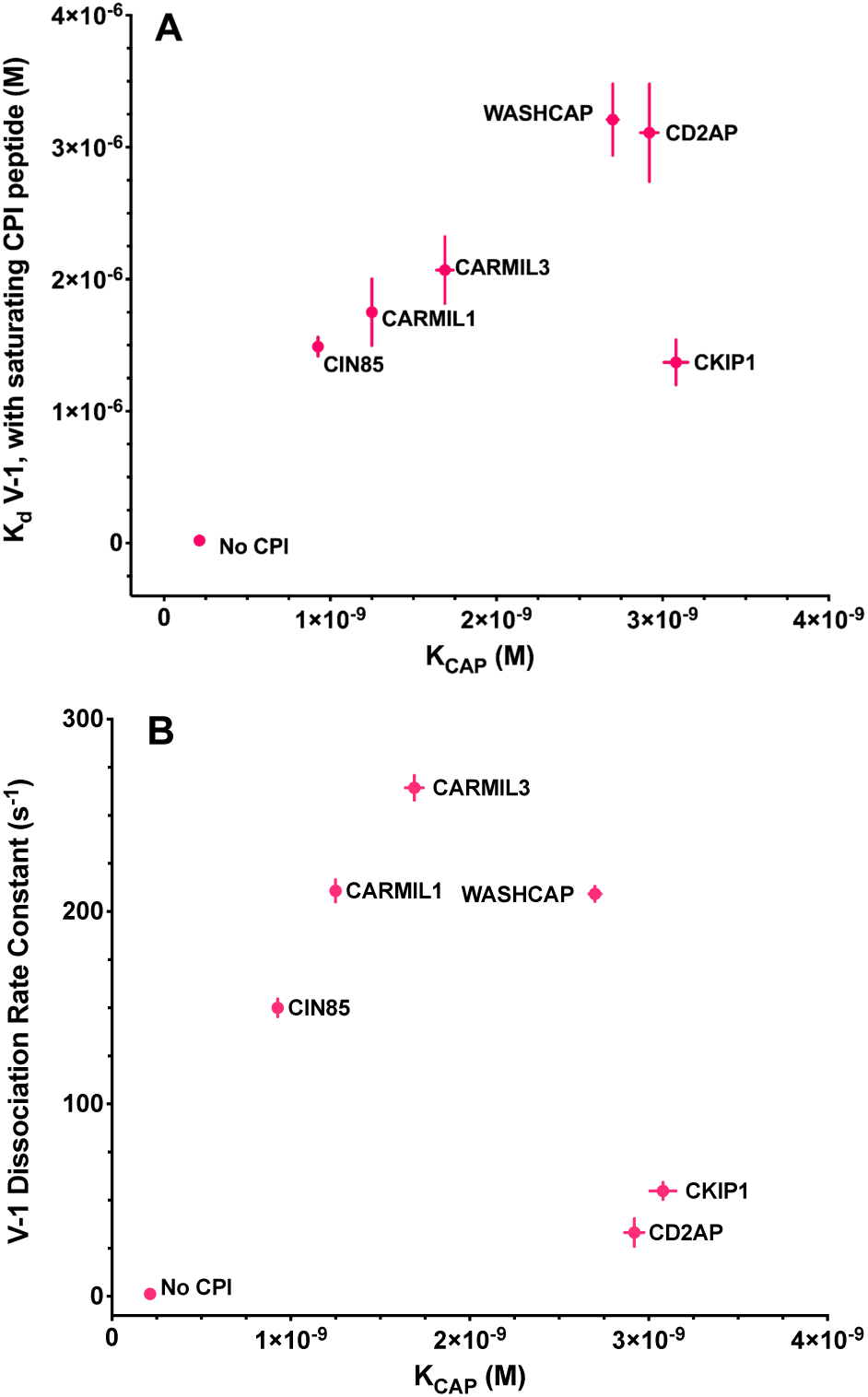
Comparison of CP:V-1 interactions with CP:actin binding affinity. Two plots comparing allosteric effects of CPI motifs on CP binding to actin with effects on the interaction of V-1 with CP. Data points are values listed in Tables 1 and 2. CPI motifs are labeled on the graph. The error bars are derived from the experimental data. When error bars are not visible, the error was less than the radius of the symbol. (A) Binding constant of V-1 for CP:CPI (CP in presence of saturating CPI) plotted versus K_CAP_, the binding constant of CP for F-actin barbed end. (B) Dissociation rate of V-1 from CP:CPI:V-1 complex plotted versus K_CAP_.

Together, these comparisons show that the different CPI motifs have distinct functional effects on CP that do not correspond simply to the binding affinity of the CPI motif for CP. Thus, the differences in amino-acid sequence in and around the CPI motif, which are conserved through evolution, appear to create differences in biochemical function.

## DISCUSSION

We found amino-acid sequence conservation among CPI-motif protein families in the regions in and around the previously defined consensus sequence for the CPI motif (*1, 2*). We asked whether these conserved amino-acid sequences would correspond with differences in biochemical functions, including the interactions of CP with V-1 and with F-actin. V-1 and F-actin have overlapping but distinct binding sites on CP, so we reasoned that such differences might exist. Indeed, our results reveal substantial differences among the CPI motifs with respect to V-1 and F-actin interactions. Our results are consistent with previous studies with a more limited scope (*1, 21*). These biochemical differences among the CPI-motif peptides may reflect differences in their cellular functions, which could be tested in the future.

We hesitate to speculate about the implications of these particular values for binding constants and rate constants, except to note that they are in the physiological range for significance relative to the concentrations of these reactants and the time scale for actin-based motility in cells. The values measured here may be valuable for mathematical models of actin filament assembly near membranes and for cell biological experiments testing the roles of CPI-motif proteins, considering situations where they function individually and in combinations.

### Thermodynamic Cycle for Interactions of CP with CPI-motif Peptides, V-1 and F-actin

To discuss the implications of the results from the different assays for the cell, it is useful to consider the binding scheme in Figure 11. Equilibrium constants for Reactions 1 through 6 were determined here experimentally, and those for Reaction 7 were calculated from the values for the other reactions.

**Figure 11.**
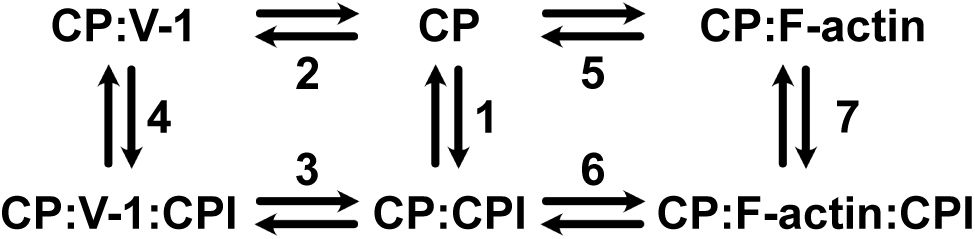
Diagram of Thermodynamic Cycle for the Binding of CP to CPI-motifs, V-1 and F-actin. K_d_s in reactions 1-4 were determined directly by experiments – fluorescence intensity and anisotropy titrations. Values for reaction 1 are in Table 1. Binding constants determined by fluorescence titrations and rate constants for dissociation of V-1 determined by stopped-flow fluorescence experiments in reactions 2 and 3 are in Table 2. K_d_s for reactions 5 and 6 (K_CAP_) determined by kinetic modelling of pyrene-actin polymerization assays are listed in Table 2. For reaction 7, K_d_s were calculated from the values for reactions 1, 5 and 6; these values are also in Table 2.

In the absence of CPI-motif peptide (top row, reactions 2 and 5, Fig. 11), K_CAP_ is 0.21 nM (Table 2, reaction 5 of Fig. 11) and the K_d_ for V-1 binding to CP is 21 nM (Table 2, reaction 2 of Fig. 11); thus, the affinity of CP for F-actin is 100 times greater than the affinity of CP for V-1. When a CPI-motif peptide is bound (i.e. in the presence of saturating CPI-motif peptide) (bottom row of Figure 11), the K_d_s for V-1 binding to CP range from 93 nM to 3.5 *µ*M (Table 2, reaction 3 of Fig. 11). Thus, the presence of a CPI-motif peptide bound to CP decreases the binding affinity of V-1 for CP by 4- to 165-fold. Values for K_CAP_ (actin capping affinity) with bound CPI-motif vary from 0.9 to 3.1 nM (Table 2, reaction 6 of Fig. 11); thus, CPI-motif peptides reduce the affinity of F-actin binding to CP by 4- to 15-fold. In the presence of CPI-motif peptides, the affinity of F-actin for CP is 450-1600 times greater than the affinity of V-1 for CP. One might then view the inhibitory sequestration of CP by V-1 as a sink mechanism, providing the cell with a pool of CP that can be activated for actin capping near a membrane by the action of a CPI motif protein.

Considering the cell further, one can envision several reactions in which a CPI-motif protein bound to a membrane might participate. A membrane-bound CPI-motif protein may encounter CP that is free (reaction 1, Fig. 11), that is bound to V-1 (reaction 4, Fig. 11) or that is bound to F-actin (reaction 7, Fig. 11). The K_d_s for CPI-motif peptides binding to CP alone vary from 8.5 to 340 nM (Table 1, reaction 1 of Fig. 11). With V-1 bound to CP, the K_d_s vary from 0.63 to 2.1 *µ*M (Table 2, reaction 4 of Fig. 11); thus, V-1 decreases the binding affinity of CPI-motif peptides to CP by 2 to 140-fold. The calculated K_d_ for CPI-motif peptides binding to CP in the presence of F-actin can be as weak as 1.3 *µ*M (Table 2, reaction 7 of Fig. 11), suggesting that high concentrations of F-actin at leading-edge cell protrusions may promote the dissociation of CP from CPI-motif proteins. In this scenario, CP that is bound to a CPI-motif protein at the membrane can be released from the membrane and cap an actin filament. The CPI-motif protein at the membranes serves as a sink mechanism, providing CP that can be used to cap barbed ends. On the other hand, the calculated K_d_ for CPI-motif peptides binding to CP in the presence of F-actin can be as strong as 61 nM (Table 2, reaction 7 of Fig. 11). In this scenario, CPI-motif protein would remove CP and uncap and actin filament.

### Consideration of CARMIL Proteins Relative to Other CPI-motif Proteins

CARMILs differ from other CPI-motif proteins in possessing a second CP-interacting region, termed the CARMIL-specific interaction (CSI) motif (*1*), just downstream of (C-terminal to) the CPI motif. This region enhances the ability of CARMILs to bind to and affect the activity of CP, relative to CPI motifs alone (*19, 35*). CARMILs also possess a third region, downstream of (C-terminal to) the CSI motif, which also affects interaction with CP and which is responsible for binding to membranes (*19, 34, 35*). Thus, while proteins with CPI motifs were found to differ rather widely in their ability to bind and affect CP, based on the results here with CPI-motif-containing peptides, we expect that CARMIL family proteins, with their CSI motifs and membrane-binding motifs, will differ from non-CARMIL CPI-motif proteins by an even larger degree. In this respect, the cellular effects and activities of CARMILs, relative to other CPI-motif proteins, may be substantially larger and more dramatic than are difference among CPI-motif proteins in general.

### Conclusions

- CPI-motifs of different protein families have amino-acid sequences conserved across evolution.
- CPI-motif peptides of different protein families bind to CP with different affinities.
- Allosteric effects of CPI motifs on binding of CP to F-actin and to V-1 also differ among CPI-motif families.
- Effects of CPI-motif peptides on the binding of CP to F-actin and V-1 show a rough correlation.
- Together, these results raise the possibility that CPI motifs have distinct functions related to the spatial and temporal regulation of actin assembly in cells.

## AUTHOR INFORMATION

### Author Contributions

The manuscript was written through contributions of all authors. All authors have given approval to the final version of the manuscript.

### Funding Sources

Funds supporting the research included: NIH 5R35GM118171 to J.A.C. and NIH 5R01GM030498 to T.M.L.

### Notes

Any additional relevant notes should be placed here.

## ACKNOWLEDGMENTS

We are grateful to Dr. Roberto Dominguez (University of Pennsylvania) for providing us with the CP:actin model in Figure 2, derived from the structures of dynactin CP and conventional actin. We are grateful to other members of the Cooper and Lohman labs for their advice and assistance and to Dr. Ken Blumer in the Department of Cell Biology & Physiology for the use of the fluorescence plate reader.

## ABBREVIATIONS

CP: Capping protein
CARMIL: Capping protein Arp2/3 Myosin-I Linker
CD2AP: CD2 associated protein
CKIP: Casein kinase 2-interacting protein
CIN85: Cbl-interacting protein of 85 kDa, aka SH3 domain-containing kinase-binding protein 1
WASH complex: Wiskott– Aldrich syndrome protein and SCAR homolog complex
FAM21: Family with sequence similarity 21 member
CapZIP: CapZ-interacting protein
V-1: myotrophin or V-1
Twf: Twinfilin
CPI-motif: Capping protein interacting motif
TCEP: tris(2-carboxyethyl)phosphine.

## SUPPORTING INFORMATION

Supplemental Figure 1, Panels A to K. Multiple sequence alignment of CPI-motif peptides. Expanded views of the results in Figure 1A, and a sequence logo for each group. Panels as follows: a, CARMIL1; b, CARMIL2; c, CARMIL3; d, CIN85; e, CD2AP; f, CapZIP; g, WASHCAP Fam21; h, CKIP-1; i, CKIP-2; j, Twinfilin-1; k, Twinfilin-2. Sequences and methods as described in the legend to Figure 1A.

Supplemental Figure 2. Phylogenetic analysis of CPI-motif peptide families. Presented as a rooted phylogenetic tree listing names of organisms.

Supplemental Figure 3. Effects of pH and salt on binding of CPI-motif peptides to CP by fluorescence anisotropy. (A) Effect of pH on binding of CARMIL1 G969-A1005 and CARMIL3 E959-M994 to CP. The negative log of K_d_ for fluorescently labeled CARMIL1 (black) and fluorescently labeled CARMIL3 (red) binding to CP are plotted versus pH. The slope for CARMIL1 CPI is 1.3 ± 0.1, and the slope for CARMIL3 CPI is 0.9 ± 0.1. The slope values indicate the net release of approximately one proton upon binding, for both peptides. Experiments were performed in 20 mM buffer, 100 mM KCl, 1 mM TCEP, 1 mM NaN_3_, 0.005 % TWEEN 20, with varying pH. pH buffers were as follows: MES pH 6.0, PIPES pH 6.5, MOPS pH 7.2, HEPES pH 7.5, EPPS pH 8.0. Competition assays with unlabeled peptides were performed for CARMIL1 at four pH values to confirm the results with fluorescently labeled peptides; the slope was 1.1 ± 0.2. (B) Effect of salt on binding of CARMIL1 and CARMIL3 to CP. The negative log of the equilibrium binding constants for fluorescent-labeled CARMIL1 (black) and fluorescent-labeled CARMIL3 (red) binding to CP are plotted versus the log of the KCl concentration. The slope for CARMIL1 CPI is −2.1 ± 0.3, and the slope for CARMIL3 CPI is −1.9 ± 0.1. These values indicate that the binding of the CPI-motif peptides to CP are accompanied by a net release of ions. Experiments were performed in 20 mM MOPS, 1 mM TCEP, 1 mM NaN_3_, 0.005 % TWEEN 20, pH 7.2, with 0.1, 0.15, 0.2, or 0.3 M KCl. Two competition experiments with unlabeled peptide were performed for CARMIL1 at 0.15 and 0.3 M KCl to confirm the result with fluorescently labeled peptides; the slope was −2.4.

Supplemental Figure 4. The effect of CP concentration on the kinetics of actin polymerization at barbed ends. Pyrene-actin fluorescence is plotted versus time. The points represent data from experiments in triplicate, and the lines are the best simultaneous (global) fits to a kinetic model for actin polymerization. CP concentrations were as follows: 0 (black), 0.75 (cyan), 2 (red), 5 (purple), 10 (magenta), 25 (green), and 50 nM (blue). K_CAP_ was 0.21 ± 0.01 nM.

## Accession IDs – UniProt

CP alpha subunit Q5RKN9

CP beta subunit Q923G3

Myotrophin (V-1) P58546

Actin P68135

CARMIL1 Q5VZK9

CARMIL3 Q8ND23

WASHCAP Q9Y4E1

CapZIP Q6JBY9

CD2AP Q9Y5K6

CIN85 Q96B97

CKIP-1 Q53GL0

Twf-1 Q91YR1

## Notes

#### Summary of Updates

Multiple minor changes in response to reviews, including data and wording.

## REFERENCES

1. Hernandez-Valladares, M., Kim, T., Kannan, B., Tung, A., Aguda, A. H., Larsson, M., Cooper, J. A., and Robinson, R. C. (2010) Structural characterization of a capping protein interaction motif defines a family of actin filament regulators., Nat Struct Mol Biol 17, 497–503.

2. Bruck, S., Huber, T. B., Ingham, R. J., Kim, K., Niederstrasser, H., Allen, P. M., Pawson, T., Cooper, J. A., and Shaw, A. S. (2006) Identification of a Novel Inhibitory Actin-capping Protein Binding Motif in CD2-associated Protein., J. Biol. Chem. 281, 19196–19203.

3. Svitkina, T. M. (2018) Ultrastructure of the actin cytoskeleton., Curr. Opin. Cell Biol. 54, 1–8.

4. Mullins, R. D., Bieling, P., and Fletcher, D. A. (2018) From solution to surface to filament: actin flux into branched networks, Biophysical Reviews 10, 1537–1551.

5. Pollard, T. D. (2016) Actin and Actin-Binding Proteins., Cold Spring Harb Perspect Biol 8, a018226.

6. Carlier, M. F., Pernier, J., Montaville, P., Shekhar, S., and Kühn, S. (2015) Control of polarized assembly of actin filaments in cell motility., Cell Mol Life Sci 72, 3051–3067.

7. Takeda, S., Minakata, S., Koike, R., Kawahata, I., Narita, A., Kitazawa, M., Ota, M., Yamakuni, T., Maeda, Y., and Nitanai, Y. (2010) Two distinct mechanisms for actin capping protein regulation--steric and allosteric inhibition., PLoS Biol. 8, e1000416.

8. Fujiwara, I., Remmert, K., Piszczek, G., and Hammer, J. A. (2014) Capping protein regulatory cycle driven by CARMIL and V-1 may promote actin network assembly at protruding edges., Proc. Natl. Acad. Sci. U S A 111, E1970–E1979.

9. Jung, G., Alexander, C. J., Wu, X. S., Piszczek, G., Chen, B. C., Betzig, E., and Hammer, J. A. (2016) V-1 regulates capping protein activity in vivo., Proc. Natl. Acad. Sci. U S A 113, E6610–E6619.

10. Cooper, J. A., and Sept, D. (2008) New insights into mechanism and regulation of actin capping protein., Int Rev Cell Mol Biol 267, 183–206.

11. Stark, B. C., Lanier, M. H., and Cooper, J. A. (2017) CARMIL family proteins as multidomain regulators of actin-based motility., Mol. Biol. Cell 28, 1713–1723.

12. Zhang, L., Tie, Y., Tian, C., Xing, G., Song, Y., Zhu, Y., Sun, Z., and He, F. (2006) CKIP-1 recruits nuclear ATM partially to the plasma membrane through interaction with ATM., Cell. Signal. 18, 1386–1395.

13. Zhao, J., Bruck, S., Cemerski, S., Zhang, L., Butler, B., Dani, A., Cooper, J. A., and Shaw, A. S. (2013) CD2AP Links Cortactin and Capping Protein at the Cell Periphery To Facilitate Formation of Lamellipodia., Mol. Cell. Biol. 33, 38–47.

14. Edwards, M., Zwolak, A., Schafer, D. A., Sept, D., Dominguez, R., and Cooper, J. A. (2014) Capping protein regulators fine-tune actin assembly dynamics., Nat. Rev. Mol. Cell Biol. 15, 677–689.

15. Edwards, M., McConnell, P., Schafer, D. A., and Cooper, J. A. (2015) CPI motif interaction is necessary for capping protein function in cells., Nat Commun 6, 8415.

16. Lanier, M. H., Kim, T., and Cooper, J. A. (2015) CARMIL2 is a novel molecular connection between vimentin and actin essential for cell migration and invadopodia formation., Mol. Biol. Cell 26, 4577–4588.

17. Edwards, M., Liang, Y., Kim, T., and Cooper, J. A. (2013) Physiological Role of the Interaction between CARMIL1 and Capping Protein., Mol. Biol. Cell 24, 3047–3055.

18. Canton, D. A., Olsten, M. E., Niederstrasser, H., Cooper, J. A., and Litchfield, D. W. (2006) The Role of CKIP-1 in Cell Morphology Depends on Its Interaction with Actin-capping Protein., J. Biol. Chem. 281, 36347–36359.

19. Zwolak, A., Uruno, T., Piszczek, G., Hammer, J. A., and Tjandra, N. (2010) Molecular basis for barbed end uncapping by CARMIL homology domain 3 of mouse CARMIL-1., J. Biol. Chem. 285, 29014–29026.

20. Johnson, B., McConnell, P., Kozlov, A. G., Mekel, M., Lohman, T. M., Gross, M. L., Amarasinghe, G. K., and Cooper, J. A. (2018) Allosteric Coupling of CARMIL and V-1 Binding to Capping Protein Revealed by Hydrogen-Deuterium Exchange., Cell Rep 23, 2795–2804.

21. Johnston, A. B., Hilton, D. M., McConnell, P., Johnson, B., Harris, M. T., Simone, A., Amarasinghe, G. K., Cooper, J. A., and Goode, B. L. (2018) A novel mode of capping protein-regulation by twinfilin., Elife 7, e41313.

22. Carlsson, A. E., Wear, M. A., and Cooper, J. A. (2004) End versus side branching by Arp2/3 complex., Biophys. J. 86, 1074–1081.

23. Anthis, N. J., and Clore, G. M. (2013) Sequence-specific determination of protein and peptide concentrations by absorbance at 205 nm., Protein Sci 22, 851–858.

24. Letunic, I., and Bork, P. (2019) Interactive Tree Of Life (iTOL) v4: recent updates and new developments., Nucleic Acids Res. 47, W256–W259.

25. Kozlov, A. G., and Lohman, T. M. (2002) Kinetic mechanism of direct transfer of Escherichia coli SSB tetramers between single-stranded DNA molecules., Biochemistry 41, 11611–11627.

26. Ramabhadran, V., Gurel, P. S., and Higgs, H. N. (2012) Mutations to the formin homology 2 domain of INF2 protein have unexpected effects on actin polymerization and severing., J. Biol. Chem. 287, 34234–34245.

27. Frieden, C. (1985) Actin and tubulin polymerization: the use of kinetic methods to determine mechanism., Annu Rev Biophys Biophys Chem 14, 189–210.

28. Cooper, J. A., and Pollard, T. D. (1985) Effect of capping protein on the kinetics of actin polymerization., Biochemistry 24, 793–799.

29. Pollard, T. D. (1986) Rate constants for the reactions of ATP- and ADP-actin with the ends of actin filaments, J. Cell Biol. 103, 2747–2754.

30. Narita, A., and Maeda, Y. (2007) Molecular determination by electron microscopy of the actin filament end structure., J. Mol. Biol. 365, 480–501.

31. Kim, T., Cooper, J. A., and Sept, D. (2010) The Interaction of Capping Protein with the Barbed End of the Actin Filament., J. Mol. Biol. 404, 794–802.

32. Zwolak, A., Fujiwara, I., Hammer, J. A., and Tjandra, N. (2010) Structural basis for capping protein sequestration by myotrophin (V-1)., J. Biol. Chem. 285, 25767–25781.

33. Narita, A., Takeda, S., Yamashita, A., and Maeda, Y. (2006) Structural basis of actin filament capping at the barbed-end: a cryo-electron microscopy study., EMBO J. 25, 5626–5633.

34. Lanier, M. H., McConnell, P., and Cooper, J. A. (2016) Cell Migration and Invadopodia Formation Require a Membrane-binding Domain of CARMIL2., J. Biol. Chem. 291, 1076–1091.

35. Kim, T., Ravilious, G. E., Sept, D., and Cooper, J. A. (2012) Mechanism for CARMIL protein inhibition of heterodimeric actin-capping protein., J. Biol. Chem. 287, 15251–15262.

