## Supplemental Figures for "Comparative analysis of CPI-motif regulation of biochemical functions of actin capping protein"

Figure S1

### a. CARMIL1

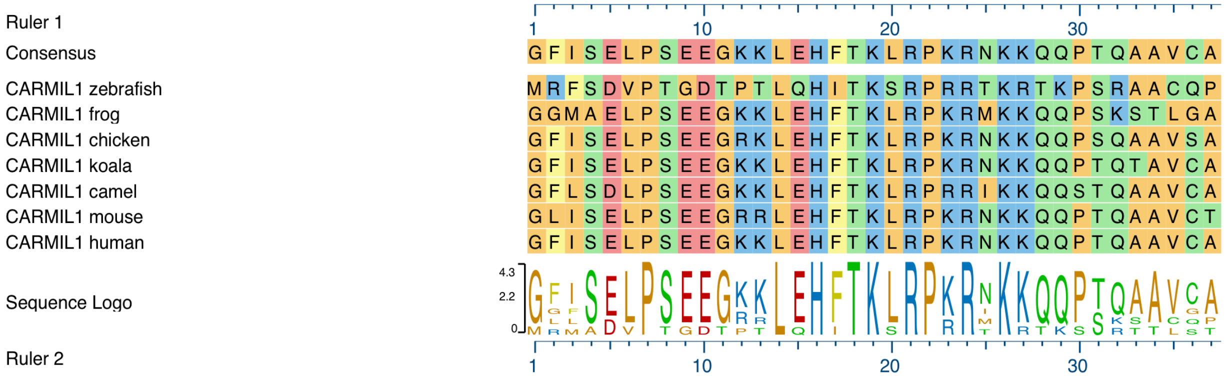


### b. CARMIL2

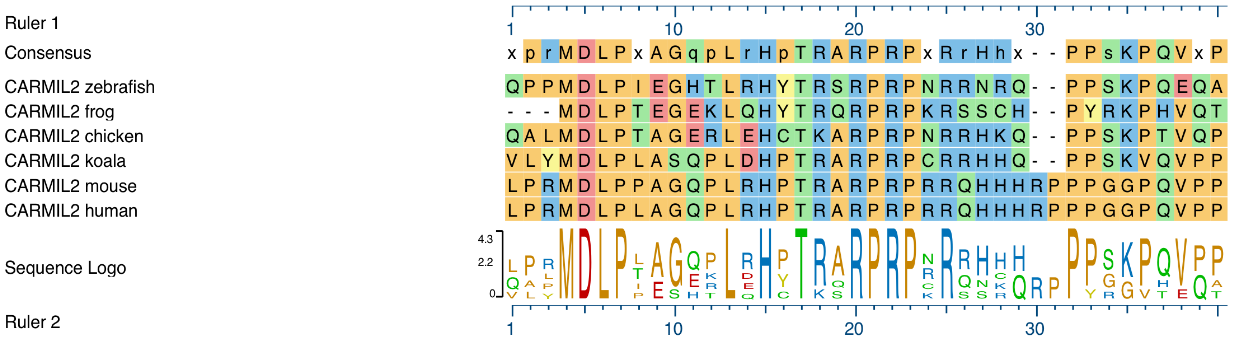


### c. CARMIL3

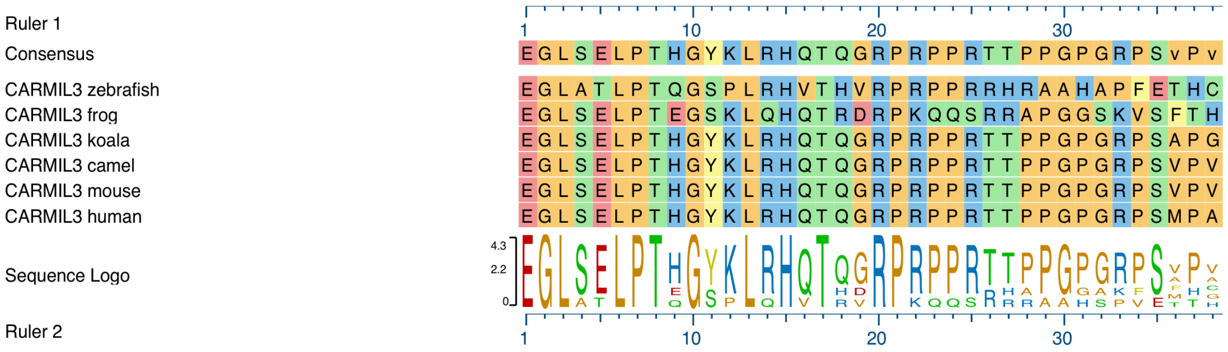

### d. CIN85

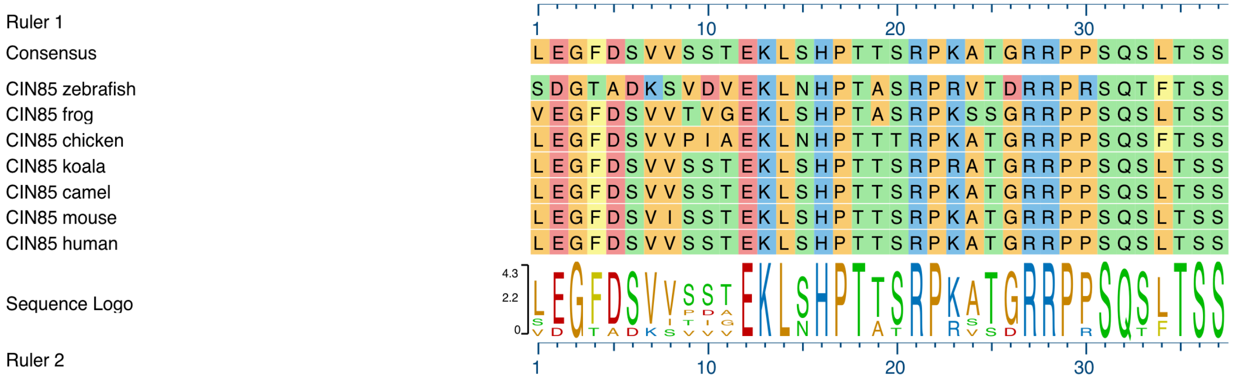


## e. CD2AP

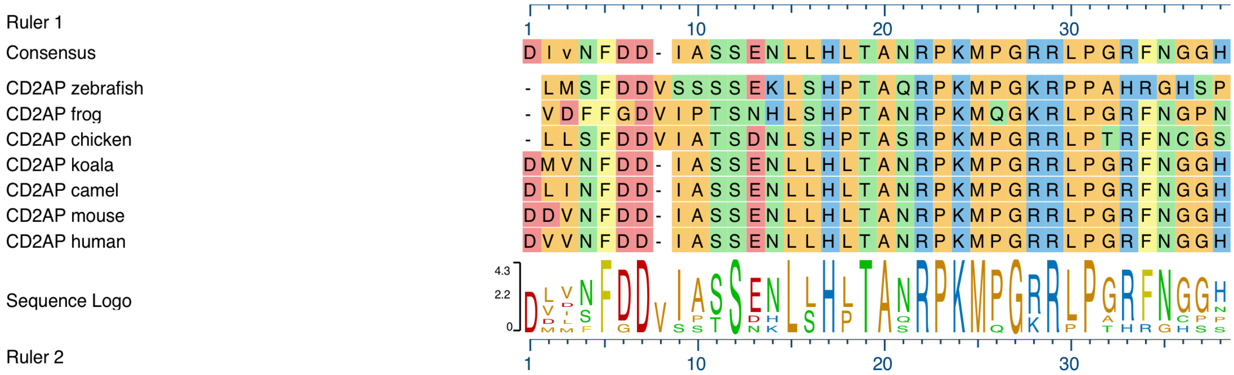


### f. CapZIP

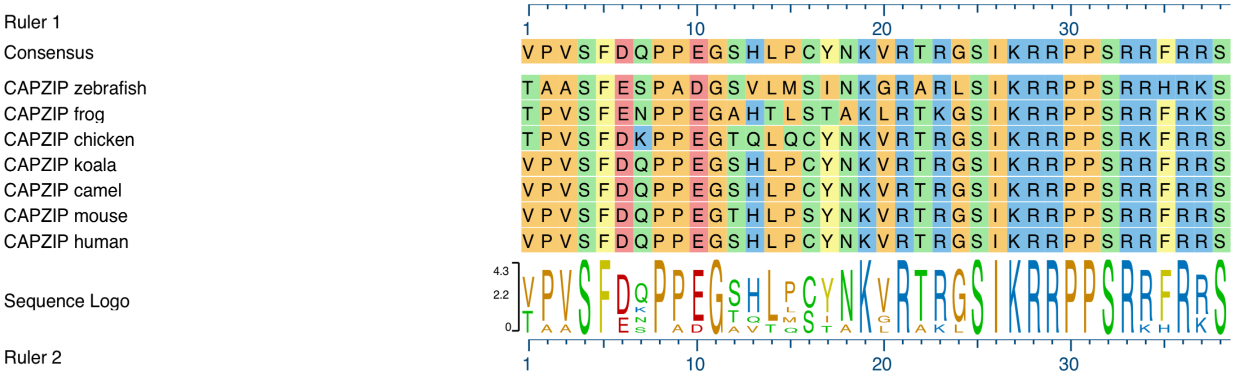


### g. WASHCAP

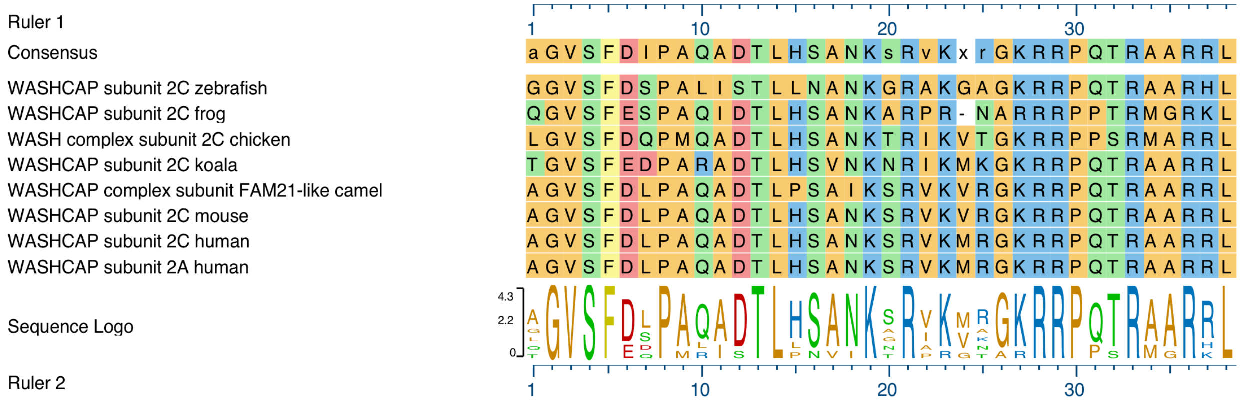

### h. CKIP-1

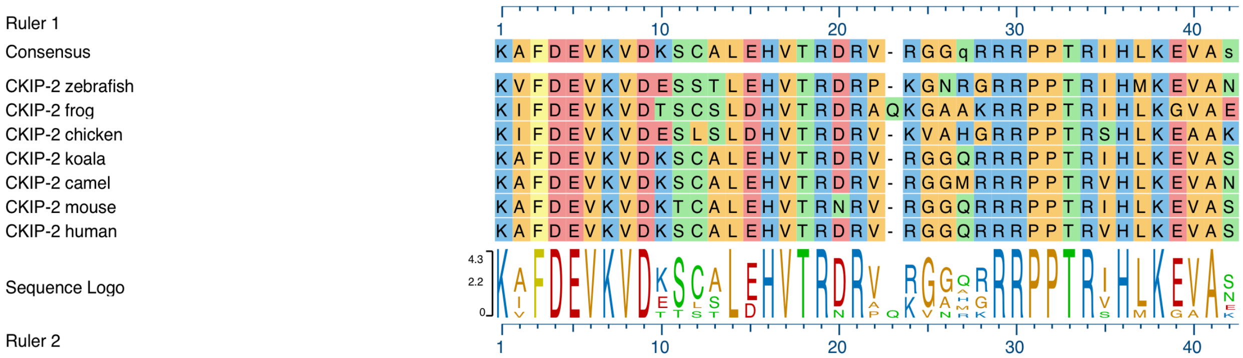


### i. CKIP-2

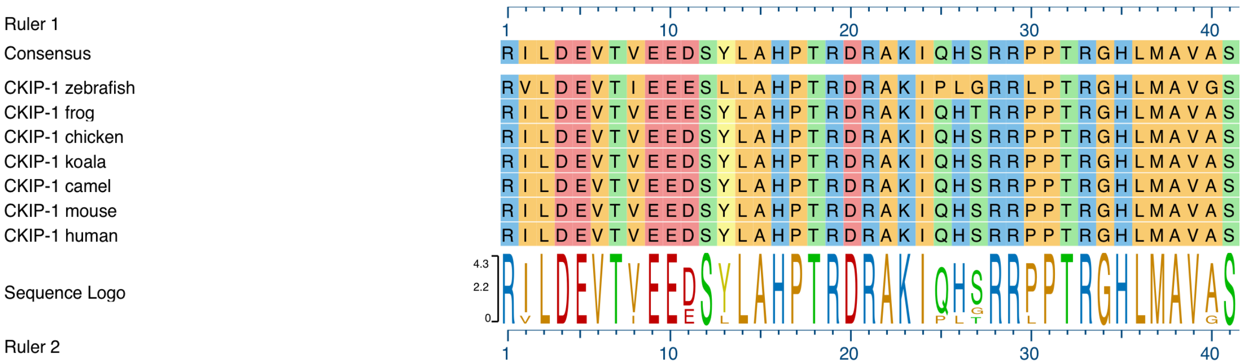


### j. Twinfilin-1

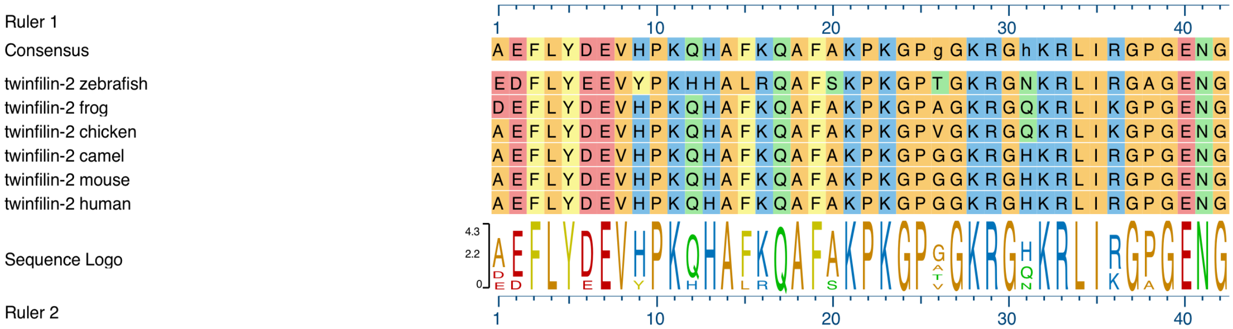


### k. Twinfilin-2

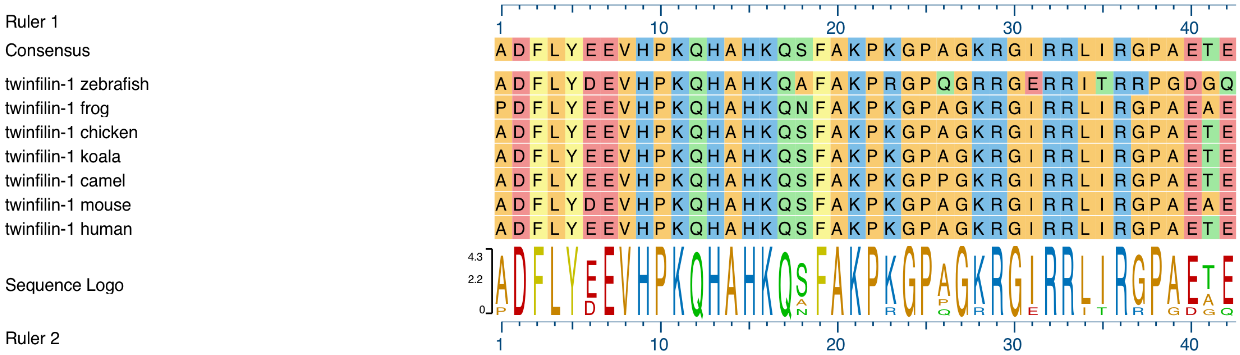

Figure S2

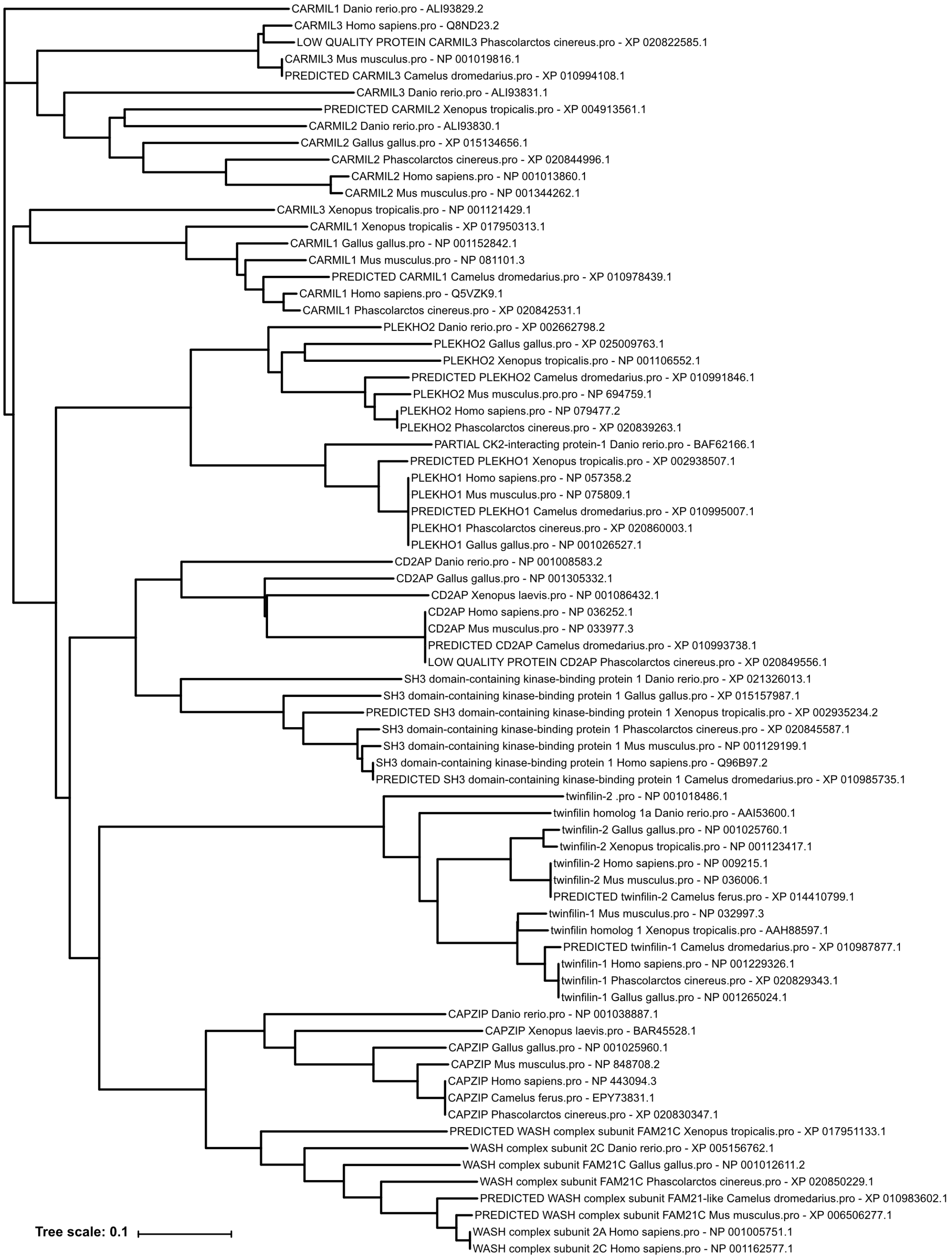
CARMIL1 Danio rerio.pro - ALI93829.2
CARMIL3 Homo sapiens.pro - Q8ND23.2
LOW QUALITY PROTEIN CARMIL3 Phascolarctos cinereus.pro - XP 020822585.1
CARMIL3 Mus musculus.pro - NP 001019816.1
PREDICTED CARMIL3 Camelus dromedarius.pro - XP 010994108.1
CARMIL3 Danio rerio.pro - ALI93831.1
PREDICTED CARMIL2 Xenopus tropicalis.pro - XP 004913561.1
CARMIL2 Danio rerio.pro - ALI93830.1
CARMIL2 Gallus gallus.pro - XP 015134656.1
CARMIL2 Phascolarctos cinereus.pro - XP 020844996.1
CARMIL2 Homo sapiens.pro - NP 001013860.1
CARMIL2 Mus musculus.pro - NP 001344262.1
CARMIL3 Xenopus tropicalis.pro - NP 001121429.1
CARMIL1 Xenopus tropicalis - XP 017950313.1
CARMIL1 Gallus gallus.pro - NP 001152842.1
CARMIL1 Mus musculus.pro - NP 081101.3
PREDICTED CARMIL1 Camelus dromedarius.pro - XP 010978439.1
CARMIL1 Homo sapiens.pro - Q5VZK9.1
CARMIL1 Phascolarctos cinereus.pro - XP 020842531.1
PLEKHO2 Danio rerio.pro - XP 002662798.2
PLEKHO2 Gallus gallus.pro - XP 025009763.1
PLEKHO2 Xenopus tropicalis.pro - NP 001106552.1
PREDICTED PLEKHO2 Camelus dromedarius.pro - XP 010991846.1
PLEKHO2 Mus musculus.pro.pro - NP 694759.1
PLEKHO2 Homo sapiens.pro - NP 079477.2
PLEKHO2 Phascolarctos cinereus.pro - XP 020839263.1
PARTIAL CK2-interacting protein-1 Danio rerio.pro - BAF62166.1
PREDICTED PLEKHO1 Xenopus tropicalis.pro - XP 002938507.1
PLEKHO1 Homo sapiens.pro - NP 057358.2
PLEKHO1 Mus musculus.pro - NP 075809.1
PREDICTED PLEKHO1 Camelus dromedarius.pro - XP 010995007.1
PLEKHO1 Phascolarctos cinereus.pro - XP 020860003.1
PLEKHO1 Gallus gallus.pro - NP 001026527.1
CD2AP Danio rerio.pro - NP 001008583.2
CD2AP Gallus gallus.pro - NP 001305332.1
CD2AP Xenopus laevis.pro - NP 001086432.1
CD2AP Homo sapiens.pro - NP 036252.1
CD2AP Mus musculus.pro - NP 033977.3
PREDICTED CD2AP Camelus dromedarius.pro - XP 010993738.1
LOW QUALITY PROTEIN CD2AP Phascolarctos cinereus.pro - XP 020849556.1
SH3 domain-containing kinase-binding protein 1 Danio rerio.pro - XP 021326013.1
SH3 domain-containing kinase-binding protein 1 Gallus gallus.pro - XP 015157987.1
PREDICTED SH3 domain-containing kinase-binding protein 1 Xenopus tropicalis.pro - XP 002935234.2
SH3 domain-containing kinase-binding protein 1 Phascolarctos cinereus.pro - XP 020845587.1
SH3 domain-containing kinase-binding protein 1 Mus musculus.pro - NP 001129199.1
SH3 domain-containing kinase-binding protein 1 Homo sapiens.pro - Q96B97.2
PREDICTED SH3 domain-containing kinase-binding protein 1 Camelus dromedarius.pro - XP 010985735.1
twinfilin-2 .pro - NP 001018486.1
twinfilin homolog 1a Danio rerio.pro - AAI53600.1
twinfilin-2 Gallus gallus.pro - NP 001025760.1
twinfilin-2 Xenopus tropicalis.pro - NP 001123417.1
twinfilin-2 Homo sapiens.pro - NP 009215.1
twinfilin-2 Mus musculus.pro - NP 036006.1
PREDICTED twinfilin-2 Camelus ferus.pro - XP 014410799.1
twinfilin-1 Mus musculus.pro - NP 032997.3
twinfilin homolog 1 Xenopus tropicalis.pro - AAH88597.1
PREDICTED twinfilin-1 Camelus dromedarius.pro - XP 010987877.1
twinfilin-1 Homo sapiens.pro - NP 001229326.1
twinfilin-1 Phascolarctos cinereus.pro - XP 020829343.1
twinfilin-1 Gallus gallus.pro - NP 001265024.1
CAPZIP Danio rerio.pro - NP 001038887.1
CAPZIP Xenopus laevis.pro - BAR45528.1
CAPZIP Gallus gallus.pro - NP 001025960.1
CAPZIP Mus musculus.pro - NP 848708.2
CAPZIP Homo sapiens.pro - NP 443094.3
CAPZIP Camelus ferus.pro - EPY73831.1
CAPZIP Phascolarctos cinereus.pro - XP 020830347.1
PREDICTED WASH complex subunit FAM21C Xenopus tropicalis.pro - XP 017951133.1
WASH complex subunit 2C Danio rerio.pro - XP 005156762.1
WASH complex subunit FAM21C Gallus gallus.pro - NP 001012611.2
WASH complex subunit FAM21C Phascolarctos cinereus.pro - XP 020850229.1
PREDICTED WASH complex subunit FAM21-like Camelus dromedarius.pro - XP 010983602.1
PREDICTED WASH complex subunit FAM21C Mus musculus.pro - XP 006506277.1
WASH complex subunit 2A Homo sapiens.pro - NP 001005751.1
WASH complex subunit 2C Homo sapiens.pro - NP 001162577.1
Tree scale: 0.1

Figure S3

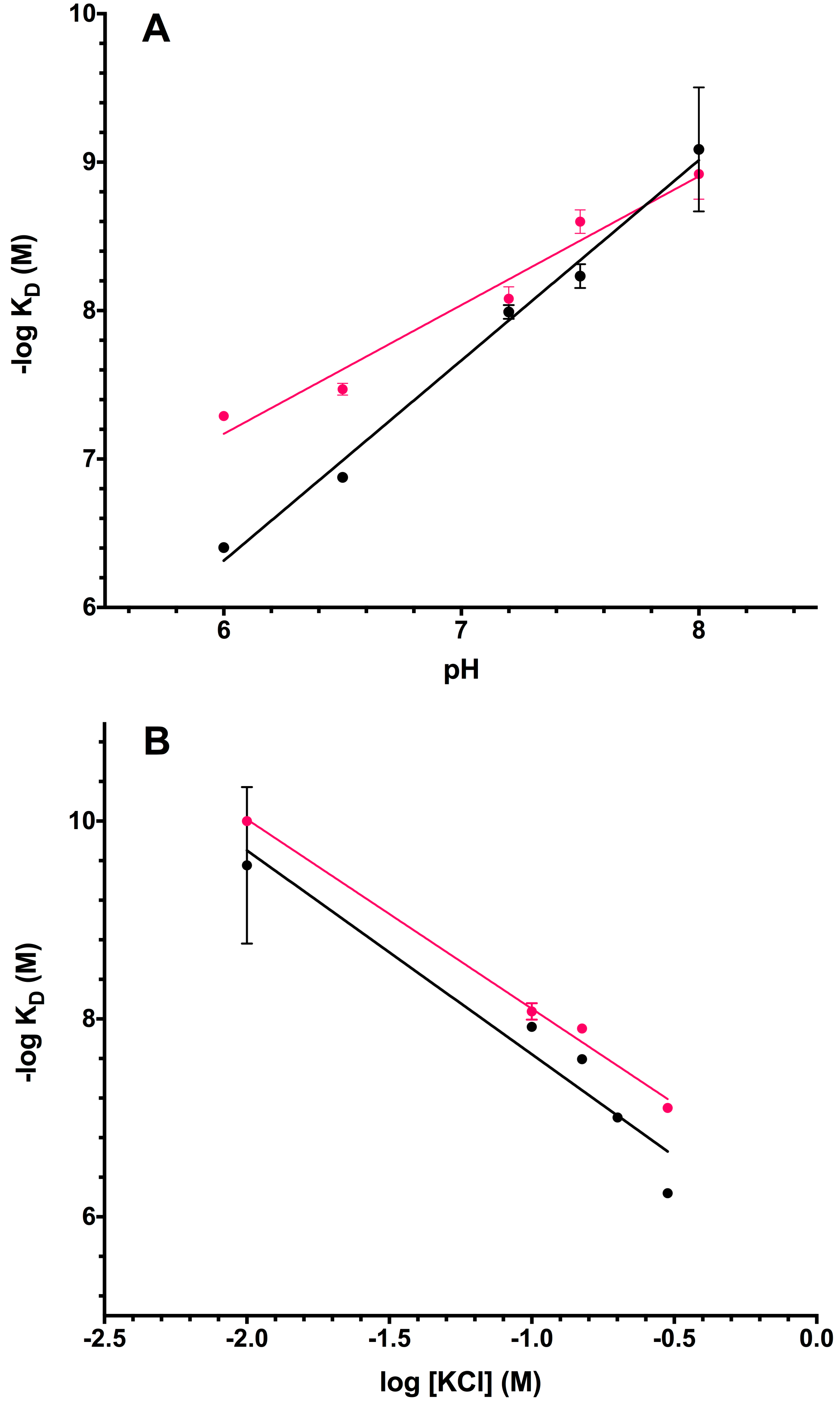

Figure S4

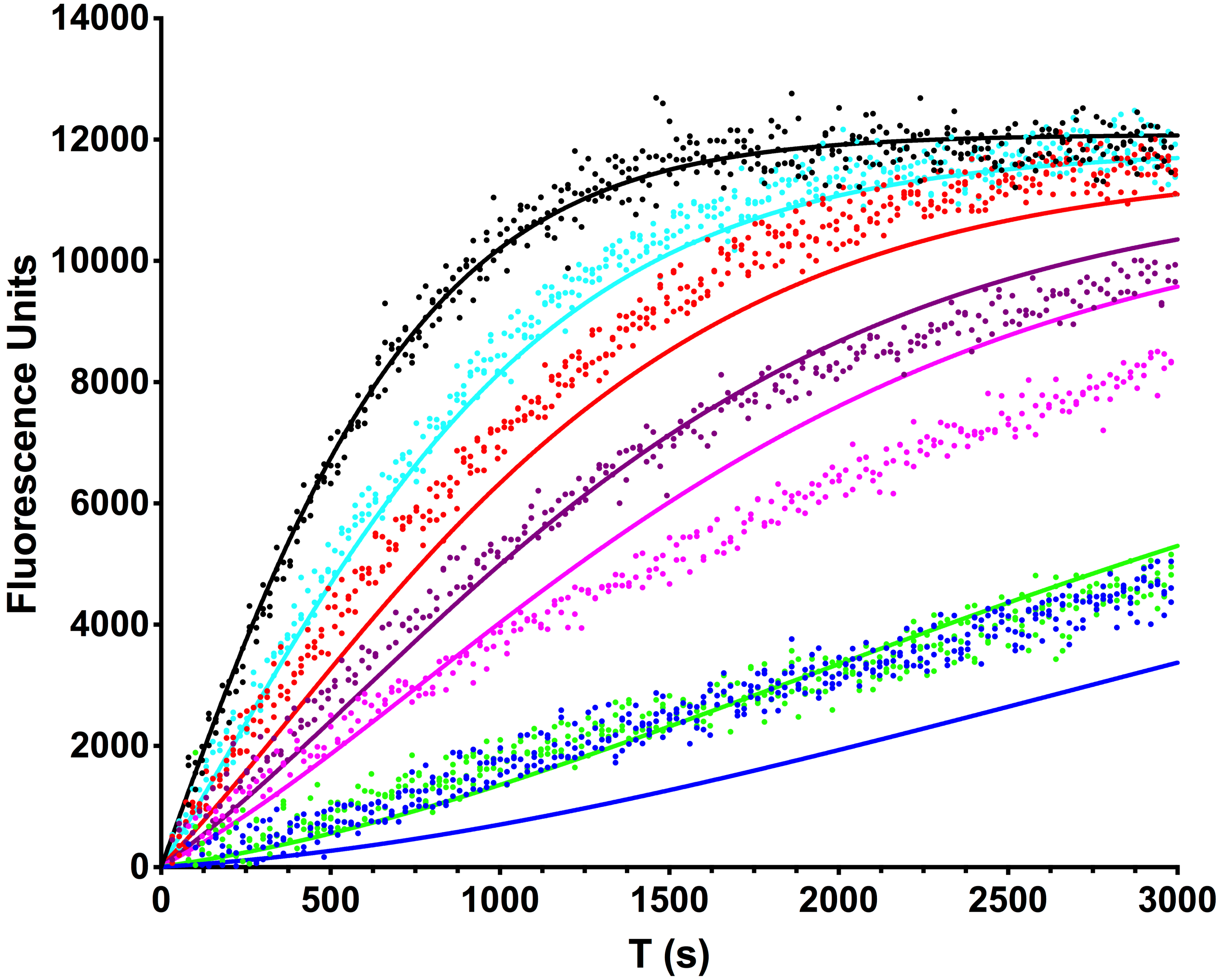

14000
12000
10000
8000
6000
4000
2000
0
Fluorescence Units
0
500
1000
1500
2000
2500
3000
T (s)
